# Genetic basis and dual adaptive role of floral pigmentation in sunflowers

**DOI:** 10.1101/2021.06.26.449999

**Authors:** Marco Todesco, Natalia Bercovich, Amy Kim, Ivana Imerovski, Gregory L. Owens, Óscar Dorado Ruiz, Srinidhi V. Holalu, Lufiani L. Madilao, Mojtaba Jahani, Jean-Sébastien Légaré, Benjamin K. Blackman, Loren H. Rieseberg

**Affiliations:** Department of Botany and Biodiversity Research Centre, University of British Columbia, Vancouver, British Columbia, Canada; Department of Biology, University of Victoria, Victoria, British Columbia, Canada; Department of Plant and Microbial Biology, University of California, Berkeley, Berkeley, California, USA; Michael Smith Laboratory and Wine Research Centre, University of British Columbia, Vancouver, British Columbia, Canada

## Abstract

Variation in floral displays, both between and within species, has been long known to be shaped by the mutualistic interactions that plants establish with their pollinators. However, increasing evidence suggests that abiotic selection pressures influence floral diversity as well. Here we analyze the genetic and environmental factors that underlie patterns of floral pigmentation in wild sunflowers. While sunflower inflorescences appear invariably yellow to the human eye, they display extreme diversity for patterns of ultraviolet pigmentation, which are visible to most pollinators. We show that this diversity is largely controlled by cis-regulatory variation at a single MYB transcription factor, HaMYB111, through accumulation of UV-absorbing flavonol glycosides. As expected, different patterns of ultraviolet pigments in flowers have a strong effect on pollinator preferences. However, variation for floral ultraviolet patterns is also associated with environmental variables, especially relative humidity, across populations of wild sunflowers. Larger ultraviolet patterns, which are found in drier environments, limit transpiration, therefore reducing water loss. The dual role of floral UV patterns in pollination attraction and abiotic responses reveals the complex adaptive balance underlying the evolution of floral traits.

## Introduction

The diversity in colour and colour patterns found in flowers is one of the most extraordinary examples of adaptive variation in the plant world. As remarkable as the variation that we can observe is, even more of it lays just outside our perception. Many species accumulate pigments that absorb ultraviolet (UV) radiation in their flowers; while these patterns are invisible to the human eye, they can be perceived by pollinators, most of which can see in the near UV (Chittka et al., 1994, Tovée, 1995). UV patterns have been shown to increase floral visibility, and to have a major influence on pollinator visitation and preference (Brock et al., 2016, Horth et al., 2014, Rae and Vamosi, 2013, Sheehan et al., 2016). Besides their role in pollinator attraction, patterns of UV-absorbing pigments in flowers have also been linked, directly or indirectly, to other biotic and abiotic factors (Gronquist et al., 2001, Koski and Ashman, 2015, Koski and Ashman, 2016).

Sunflowers have come to enjoy an iconic status in popular culture (as testified by the, arguably dubious, honour of being one of the only five flower species with a dedicated emoji (Unicode.org, 2020)). This is despite being apparently largely immune from the aforementioned diversification for flower colour; all 50 species of wild sunflowers have ligules (the enlarged modified petals of the outermost whorl of florets in the sunflower inflorescences) that appear of the same bright yellow colour to the human eye. However, ligules also accumulate UV-absorbing pigments at their base, while their tip reflects UV radiation (Harborne and Smith, 1978, Wojtaszek and Maier, 2014). Across the whole inflorescence, this results in a bullseye pattern with an external UV-reflecting ring and an internal UV-absorbing ring. Besides their well-described role in pollinator attraction, UV bullseyes have been proposed to act as nectar guides, helping pollinators orient towards nectar rewards once they land on the petal, although recent experiments have challenged this hypothesis (Koski and Ashman, 2014). Considerable variation in the size of UV bullseye patterns has been observed between and within plant species (Koski and Ashman, 2013, Koski and Ashman, 2016); however, little is known about the ecological factors that drive this variation, or the genetic determinants that control it.

## Results and Discussion

### Floral UV patterns in wild sunflowers

A preliminary screening of 15 species of wild sunflowers, as well as cultivated sunflower, suggested that UV bullseye patterns are common across sunflower species (Figure 1 – figure supplement 1). We also observed substantial within-species variation for the size of UV floral patterns. Variation for UV bullseye size was previously reported in the silverleaf sunflower *Helianthus argophyllus*; however, genetic mapping resolution was insufficient to identify individual causal genes (Moyers et al., 2017). To better understand the function and genetic regulation of variation for floral UV pigmentation, we focused on two widespread species of annual sunflowers, *H. annuus* (the progenitor of the domesticated sunflower) and *H. petiolaris*; both have broad distributions across North America, but the latter prefers sandier soils (Heiser and Smith, 1969, Todesco et al., 2020). Over two growing seasons, we measured UV floral patterns (as the proportion of the ligule that absorbs UV radiation, henceforth “Ligule UV proportion” or “LUVp”) in 1589 *H. annuus* individuals derived from 110 distinct natural populations, and 351 *H. petiolaris* individuals from 40 populations, grown in common garden experiments in Vancouver, Canada (Todesco et al., 2020) (Figure 1a,b; Figure 1 – source data 1). While extensive variation was observed within both species, it was particularly striking for *H. annuus*, which displayed a phenotypic continuum from ligules with almost no UV pigmentation to ligules that were entirely UV-absorbing (Figure 1c-e; Figure 1 – source data 2). A relatively high proportion of *H. annuus* individuals (∼13%) had completely UV-absorbing ligules and therefore lacked UV “nectar guides”, suggesting that pollinator orientation is not a necessary function of floral UV pigmentation in sunflower.

**Figure 1:**
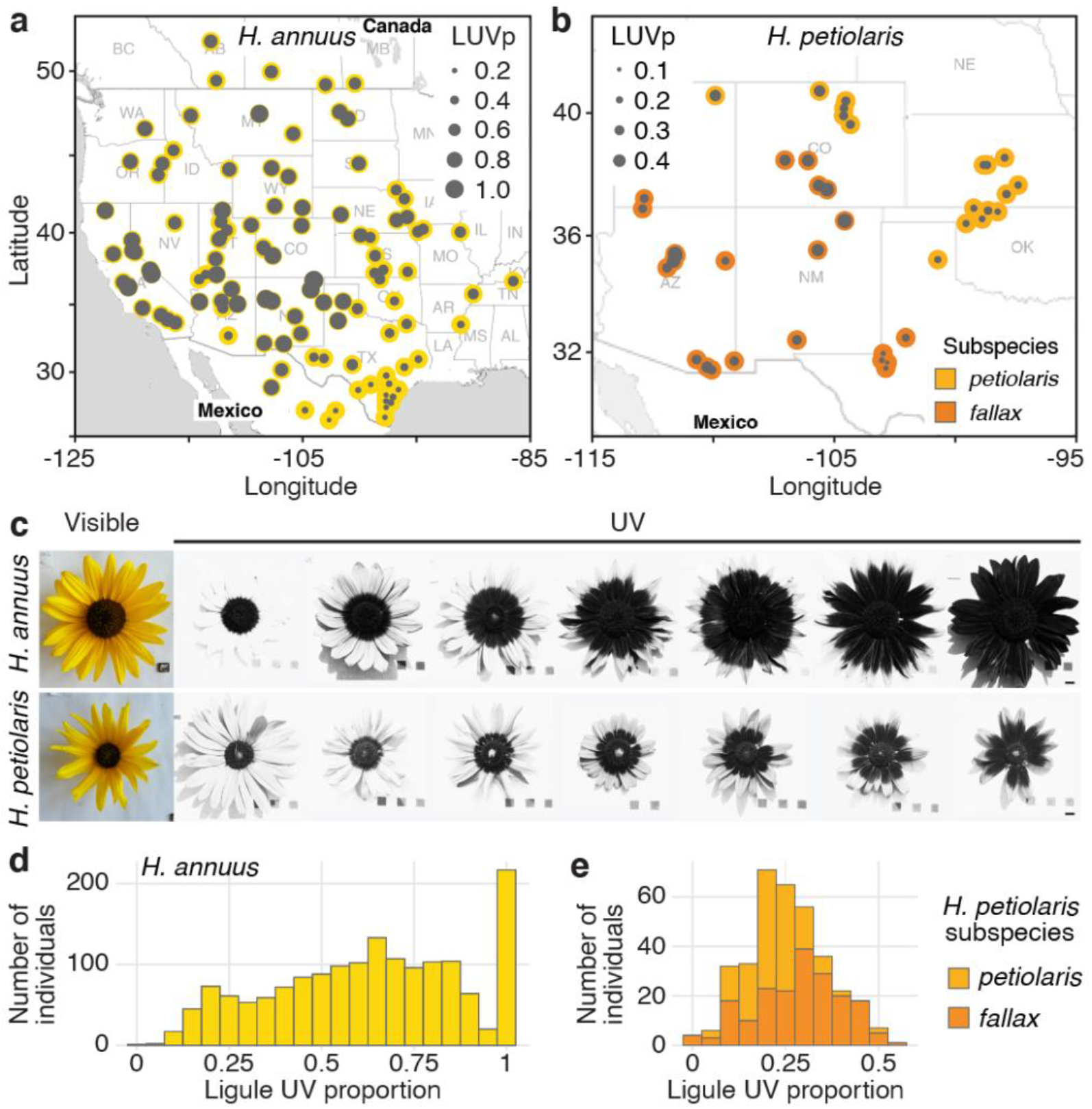
Diversity for floral UV pigmentation patterns in wild sunflowers. **a,** Geographic distribution of sampled populations for *H. annuus* and **b,** *H. petiolaris*. Yellow/orange dots represent different populations, overlaid grey dots size is proportional to the population mean LUVp. **c,** Range of variation for floral UV pigmentation patterns in the two species. Scale bar = 1 cm. **d,** LUVp values distribution for *H. annuus* and **e,** *H. petiolaris* subspecies.

### Genetic control of floral UV patterning

To identify the loci controlling variation for floral UV patterning, we performed a genome-wide association study (GWAS). We used a subset of the phenotyped plants (563 of the *H. annuus* and all 351 *H. petiolaris* individuals) for which we previously generated genotypic data at >4.6M high-quality single-nucleotide polymorphisms (SNPs) (Todesco et al., 2020). Given their relatively high level of genetic differentiation, analyses were performed separately for the *petiolaris* and *fallax* subspecies of *H. petiolaris* (Todesco et al., 2020). We detected several genomic regions significantly associated with UV patterning in *H. petiolaris petiolaris*, and a particularly strong association (*P* = 5.81e^-25^) on chromosome 15 in *H. annuus* (Figure 2a,b; Figure 2 – figure supplement 1). The chromosome 15 SNP with the strongest association with ligule UV pigmentation patterns in *H. annuus* (henceforth “Chr15_LUVp SNP”) explained 62% of the observed variation, and allelic distributions at this SNP closely matched that of floral UV patterns (Figure 2c, compare to Figure 1a; Figure 1 – source data 2).

**Figure 2:**
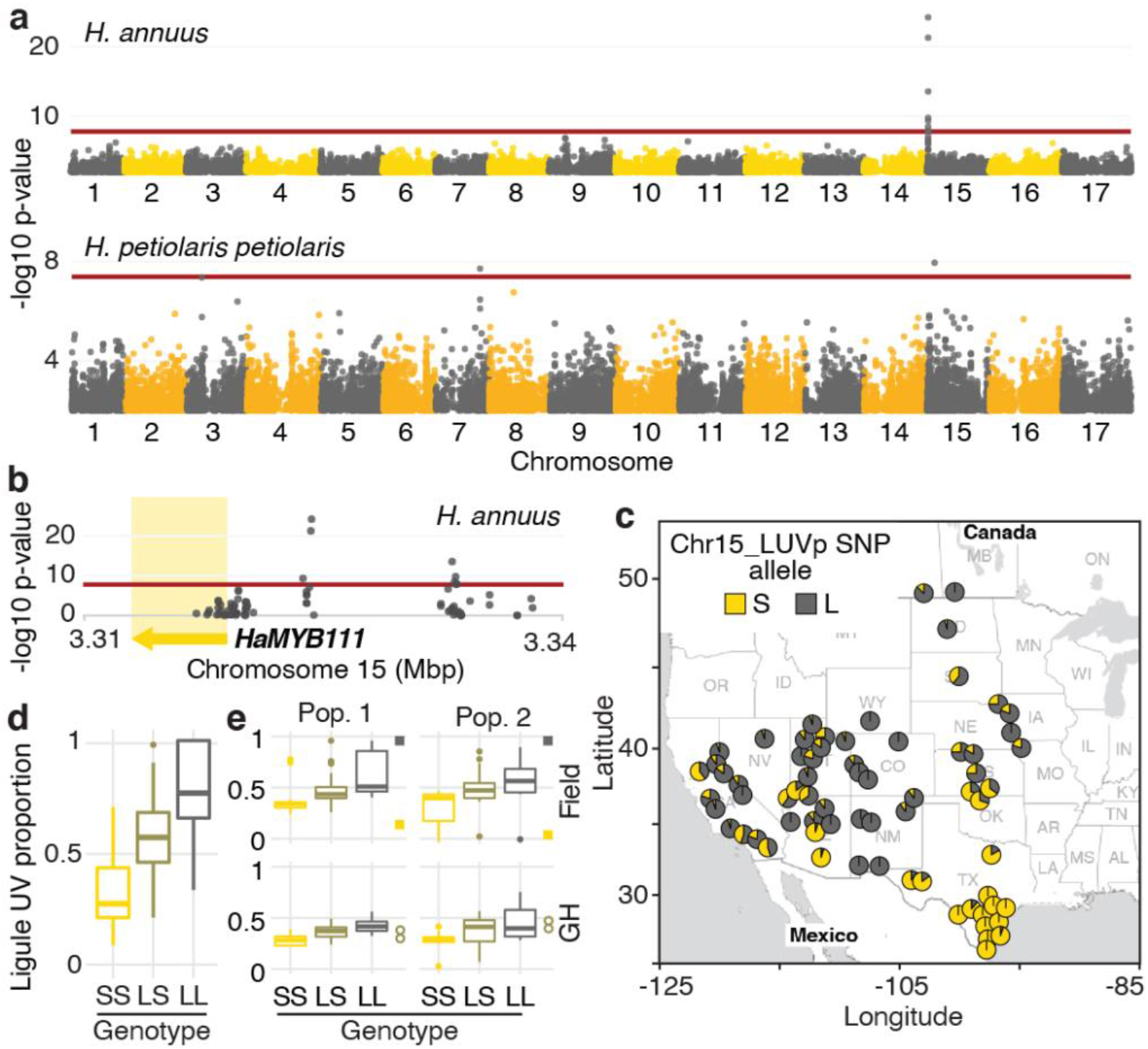
A single locus explains most of the variation in floral UV patterning in *H. annuus.* **a,** LUVp GWAS. **b,** Zoomed-in Manhattan plot for the chromosome 15 LUVp peak in *H. annuus*. Red lines represent 5% Bonferroni-corrected significance. GWAs were calculated using two-sided mixed models. Number of individuals: *n* = 563 individuals (*H. annuus*); n = 159 individuals (*H. petiolaris petiolaris*). Only positions with - log10 p-value >2 are plotted. **c,** Geographic distribution of Chr15_LUVp SNP allele frequencies in *H. annuus*. L = Large and S = Small allele. **d,** LUVp associated with different genotypes at Chr15_LUVp SNP in natural populations of *H. annuus* grown in a common garden. All pairwise comparisons are significant for *P* < 10^-16^ (one-way ANOVA with post-hoc Tukey HSD test, *F* = 438, df = 2; *n* = 563 individuals). LUVp values and genotype data for Chr15_LUVp SNP are reported in Figure 1 – source data 2. **e,** LUVp associated with different genotypes at Chr15_LUVp SNP in *H. annuus* F_2_ populations grown in the field or in a greenhouse (GH). Measurements for the parental generations are shown: squares = grandparents (field-grown); empty circles = F1 parents (greenhouse-grown; figure 2 – figure supplement 2). Boxplots show the median, box edges represent the 25^th^ and 75^th^ percentiles, whiskers represent the maximum/minimum data points within 1.5x interquartile range outside box edges. Differences between genotypic groups are significant for *P* = 0.0057 (Pop. 1 Field, one-way ANOVA, *F* = 5.73, df = 2; *n* = 54 individuals); *P* = 0.0021 (Pop. 2 Field, one-way ANOVA, *F* = 7.02, df = 2; *n* = 50 individuals); *P* = 0.00015 (Pop. 1 GH, one-way ANOVA, *F* = 11.13, df = 2; *n* = 42 individuals); *P* = 0.054 (Pop. 2 GH, one-way ANOVA, *F* = 3.17, df = 2; *n* = 38 individuals). P-values for pairwise comparisons for panels d and e are reported in the source data for this figure.

Genotype at the Chr15_LUVp SNP had a remarkably strong effect on the size of UV bullseyes in inflorescences. Individuals homozygous for the “large” (L) allele had a mean LUVp of 0.78 (st.dev ±0.16), meaning that ∼3/4 of the ligule was UV-absorbing, while individuals homozygous for the “small” (S) allele had a mean LUVp of 0.33 (st.dev. ±0.15), meaning that only the basal ∼1/3 of the ligule absorbed UV radiation. Consistent with the trimodal LUVp distribution observed for *H. annuus* (Figure 1d), alleles at this locus showed additive effects, with heterozygous individuals having intermediate phenotypes (LUVp = 0.59 ± 0.18; Figure 2d). The association between floral UV patterns and the Chr15_LUVp SNP was confirmed in the F_2_ progeny of crosses between plants homozygous for the L allele (with completely UV-absorbing ligules; LUVp = 1) and for the S allele (with a small UV-absorbing patch at the ligule base; LUVp < 0.18; Figure 2e; Figure 2 – figure supplement 2). Average LUVp values were lower, and their range smaller, when these populations were grown in a greenhouse rather than in a field. Plants in the greenhouse experienced relatively uniform temperatures and humidity, and were shielded from most UV radiation. This suggests that while floral UV patterns have a strong genetic basis (consistent with previous observations (Koski and Ashman, 2013)), their expression is also affected by the environment.

### *HaMYB111* regulates UV pigment production

While no obvious candidate genes were found for the GWA peaks for floral UV pigmentation in *H. petiolaris petiolaris*, the *H. annuus* chromosome 15 peak is ∼5 kbp upstream of *HaMYB111*, a sunflower homolog of the *Arabidopsis thaliana AtMYB111* gene (Figure 2b). Together with AtMYB11 and AtMYB12, AtMYB111 is part of a small family of transcription factors (also called PRODUCTION OF FLAVONOL GLYCOSIDES, PFG) that controls the expression of genes involved in the production of flavonol glycosides in Arabidopsis (Stracke et al., 2007). Flavonol glycosides are a subgroup of flavonoids known to fulfill a variety of functions in plants, including protection against abiotic and biotic stresses (e.g. UV radiation, cold, drought, herbivory) (Pollastri and Tattini, 2011). Crucially, they absorb strongly in the near UV range (300-400 nm), and are the pigments responsible for floral UV patterns in several plant species (Brock et al., 2016, Rieseberg and Schilling, 1985, Sheehan et al., 2016, Thompson et al., 1972). For instance, alleles of a homolog of *AtMYB111* are responsible for the evolutionary gain and subsequent loss of flavonol accumulation and UV absorption in flowers of *Petunia* species, associated with two successive switches in pollinator preferences (from bees, to hawkmoths, to hummingbirds (Sheehan et al., 2016)). A homolog of *AtMYB12* has also been associated with variation in floral UV patterns in *Brassica rapa* (Brock et al., 2016). Consistent with this, we found flavonol glycosides to be the main UV-absorbing pigments in sunflower ligules, accumulating at much higher levels at their base, and in ligules of plants with larger LUVp (Figure 3a,b).

**Figure 3:**
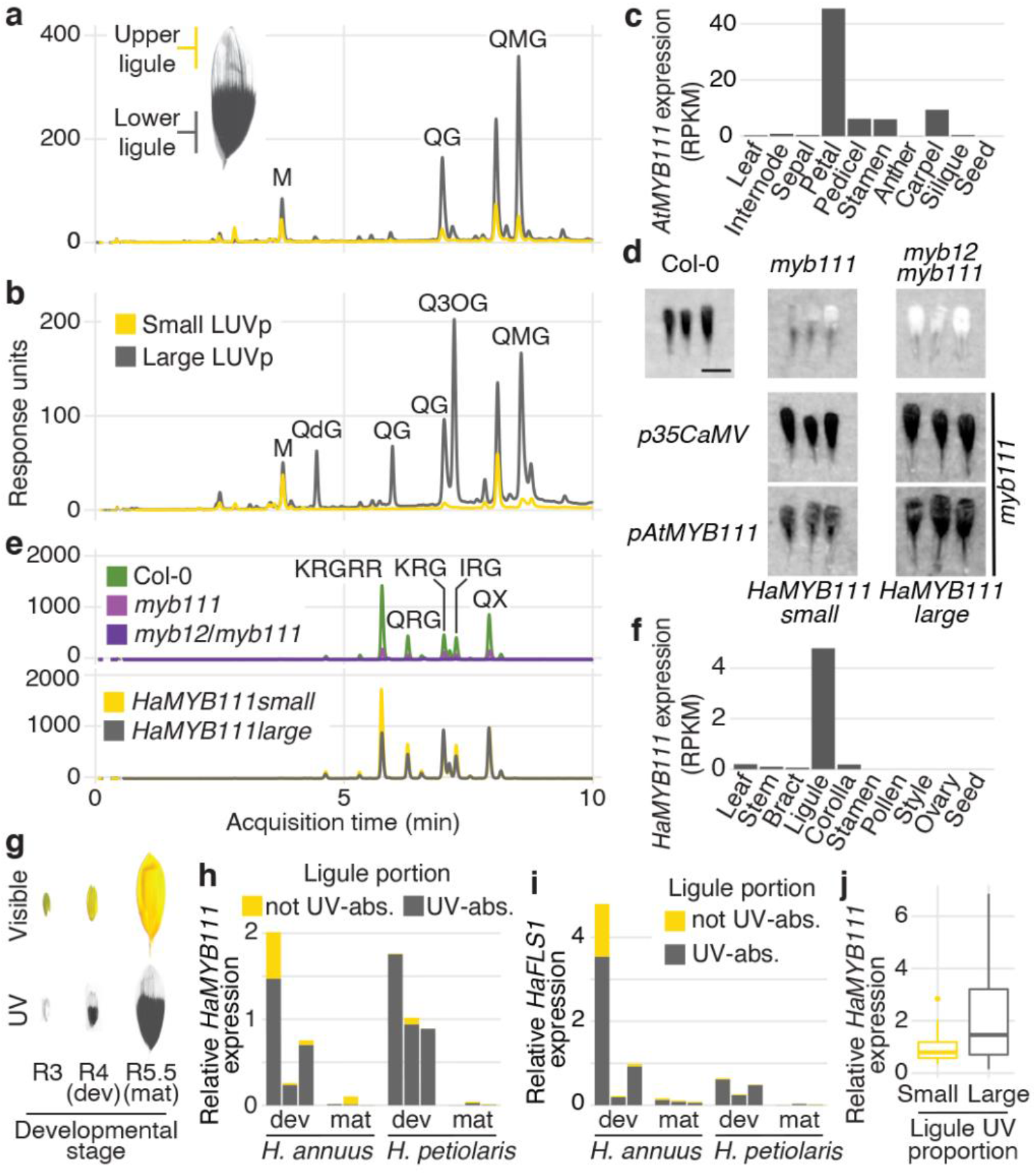
*MYB111* is associated with floral UV pigmentation patterns and flavonol accumulation in sunflower and Arabidopsis. **a,** UV chromatograms (350 nm) for methanolic extracts of the upper and lower third of ligules with intermediate UV patterns, and **b,** of ligules with large and small floral UV patterns. Peaks corresponding to flavonols are labelled (Figure 3 – source data 1). **c,** Expression levels of *AtMYB111* in Arabidopsis. RPKM = Reads Per Kilobase of transcript per Million mapped reads. **d,** UV pictures of Arabidopsis petals. *HaMYB111* from *H. annuus* plants with small or large LUVp was introduced into the Arabidopsis *myb111* mutant under the control of a constitutive promoter (*p35SCaMV*) or of the promoter of the Arabidopsis homolog (*pAtMYB111*). All petals are white in the visible range (Figure 3 – figure supplement 1). Scale bar = 1mm. **e,** UV chromatograms (350 nm) for methanolic extracts of petals of Arabidopsis lines. Upper panel: wild-type Col-0 and mutants. Bottom panel: *p35SCaMV::HaMYB111* lines in *myb111* background. Peaks corresponding to flavonols are labelled (Figure 3 – source data 1). **f,** Expression levels of *HaMYB111* in cultivated sunflower. **g,** Pigmentation patterns in ligules of wild *H. annuus* at different developmental stages: R3 = closed inflorescence bud; R4 = inflorescence bud opening; R5 = inflorescence fully opened. **h,**Expression levels in the UV-absorbing (grey) and UV-reflecting (yellow) portion of mature (mat) and developing (dev) ligules for *HaMYB111* and **i,** *HaFLS1*, one of its putative targets. Each bar represents a biological replicate (different inflorescence from a same individual). **j,** *HaMYB111* expression levels in field-grown wild *H. annuus* with different floral UV pigmentation patterns. The difference between the two groups is significant for *P* = 0.009 (Welch t-test, *t* = 2.81, df = 27.32, two-sided; *n* = 46 individuals). Boxplots show the median, box edges represent the 25^th^ and 75^th^ percentiles, whiskers represent the maximum/minimum data points within 1.5x interquartile range outside box edges.

*AtMYB12* and *AtMYB111* are known to have the strongest effect on flavonol glycoside accumulation in Arabidopsis (Stracke et al., 2007, Stracke et al., 2010). We noticed, from existing RNAseq data, that *AtMYB111* expression levels are particularly high in petals (Klepikova et al., 2016) (Figure 3c), and found that Arabidopsis petals, while uniformly white in the visible spectrum, absorb strongly in the UV (Figure 3d; Figure 3 – figure supplement 1). To our knowledge, this is the first report of floral UV pigmentation in Arabidopsis, a highly selfing species which is seldom insect-pollinated (Hoffmann et al., 2003). Accumulation of flavonol glycosides in petals is strongly reduced, and UV pigmentation is almost completely absent, in petals of mutants for *AtMYB111* (*myb111*; Figure 3d,e). UV absorbance is further reduced in petals of double mutants for *AtMYB12* and *AtMYB111* (*myb12/111*). However, petals of the single mutant for *AtMYB12* (*myb12*), which is expressed at low levels throughout the plant (Klepikova et al., 2016), are indistinguishable from wild-type plants (Figure 2 – figure supplement 1). This shows that flavonol glycosides are responsible for floral UV pigmentation also in Arabidopsis, and that *AtMYB111* plays a fundamental role in controlling their accumulation in petals.

To confirm that sunflower *HaMYB111* is functionally equivalent to its Arabidopsis homolog, we introduced it into *myb111* plants. Expression of *HaMYB111*, either under the control of a constitutive promoter or of the endogenous *AtMYB111* promoter, restored petal UV pigmentation and induced accumulation of flavonol glycosides (Figure 3d,e). *HaMYB111* coding sequences obtained from wild sunflowers with large or small LUVp were equally effective at complementing the *myb111* mutant. Together with the observation that the strongest GWAS association with LUVp fell in the promoter region of *HaMYB111*, these results suggest that differences in the effect of the “small” and “large” alleles of this gene on floral UV pigmentation are not due to differences in protein function, but rather to differences in gene expression.

Analysis of *HaMYB111* expression in cultivated sunflower revealed that, consistent with a role in floral UV pigmentation and similar to its Arabidopsis counterpart, it is expressed specifically in ligules, and it is almost undetectable in other tissues (Badouin et al., 2017) (Figure 3f). Similar to observations in *Rudbeckia hirta*, another member of the *Heliantheae* tribe (Schlangen et al., 2009), UV pigmentation is established early in ligule development in both *H. annuus* and *H. petiolaris*, as their visible colour turns from green to yellow before the inflorescence opens (R4 developmental stage (Schneiter and Miller, 1981); Figure 3g; Figure 3 – figure supplement 2). *HaMYB111* is highly expressed in the part of the ligule that accumulates UV-absorbing pigments, and especially in developing ligules, consistent with a role in establishing pigmentation patterns (Figure 3h). We also observed a matching expression pattern for *HaFLS1*, the sunflower homolog of a gene encoding one of the main enzymes controlling flavonol biosynthesis in Arabidopsis (*FLAVONOL SYNTHASE 1, AtFLS1*), whose expression is regulated directly by *AtMYB111* (Stracke et al., 2007) (Figure 3i). Finally, we compared *HaMYB111* expression levels in a set of 46 field-grown individuals with contrasting LUVp values, representing 21 different wild populations. *HaMYB111* expression levels differed significantly between the two groups (*P* = 0.009; Figure 3j). Variation in expression levels within phenotypic classes was quite large; this is likely due in part to the strong dependence of *HaMYB111* expression on developmental stage (Figure 3g), and the difficulty of accurately establishing matching ligule developmental stages across diverse wild sunflowers.

These expression analyses further point to *cis*-regulatory rather than coding sequence differences between *HaMYB111* alleles being responsible for LUVp variation. Accordingly, direct sequencing of the *HaMYB111* locus from multiple wild *H. annuus* individuals, using a combination of Sanger sequencing and long PacBio HiFi reads, identified no coding sequence variants associated with differences in floral UV patterns, or with alleles at the Chr15_LUVp SNP. However, we observed extensive variation in the promoter region of *HaMYB111*, differentiating wild *H. annuus* alleles from each other and from the reference assembly for cultivated sunflower. Relaxing quality filters to include less well-supported SNPs in our LUVp GWAS did not identify additional variants with stronger associations than Chr15_LUVp SNP (Figure 2 – figure supplement 2). However, many of the polymorphisms we identified by direct sequencing were either larger insertions/deletions (indels) or fell in regions that were too repetitive to allow accurate mapping of short reads, and would not be included even in the expanded SNP dataset. While several of these variants in the promoter region of *HaMYB111* appeared to be associated with the Chr15_LUVp SNP, further studies will be required to confirm this, and to identify their eventual effects on *HaMYB111* activity.

Interestingly, when we sequenced the promoter region of *HaMYB111* in several *H. argophyllus* and *H. petiolaris* individuals, we found that they all carried the S allele at the Chr15_LUVp SNP. Similarly, in a set of previously re-sequenced wild sunflowers, we found the S allele to be fixed in several perennial (*H. decapetalus*, *H. divaricatus* and *H. grosseserratus*) and annual sunflower species (*H. argophyllus*, *H. niveus*, *H. debilis*), and to be at >0.98 frequency in *H. petiolaris*, suggesting that the “small” haplotype is ancestral. Conversely, the L allele at Chr15_LUVp SNP was almost fixed (>0.98 frequency) in a set of 285 cultivated sunflower lines (Mandel et al., 2013). Consistent with these patterns, UV bullseyes are considerably smaller in *H. argophyllus* (mean LUVp ± st.dev. = 0.27 ± 0.09), *H. niveus* (0.15 ± 0.09), and *H. petiolaris* (0.27 ± 0.12; Figure 1e) than in cultivated sunflower lines (0.62 ± 0.23). Additionally, while 50 of the cultivated sunflower lines had completely or almost completely UV-absorbing ligules (LUVp > 0.8), no such case was observed in the other three species (Figure 1 – figure supplement 2).

### A dual role for floral UV pigmentation

Although our results show that *HaMYB111* explains most of the variation in floral UV pigmentation patterns in wild *H. annuus*, why such variation exists in the first place is less clear. Several hypotheses have been advanced to explain the presence of floral UV patterns and their variability. Like their visible counterparts, UV pigments play a fundamental role in pollinator attraction (Horth et al., 2014, Koski and Ashman, 2014, Rae and Vamosi, 2013, Sheehan et al., 2016). For example, in *Rudbeckia* species, artificially increasing the size of bullseye patterns to up to 90% of the petal surface resulted in rates of pollinator visitation equal to or higher than wild-type flowers (which have on average 40-60% of the petal being UV-absorbing). Conversely, reducing the size of the UV bullseye had a strong negative effect on pollinator visitation (Horth et al., 2014). To test whether the relative size of UV bullseye patterns affected pollination, we compared insect visitation rates for wild *H. annuus* lines with contrasting UV bullseye patterns. An initial experiment, carried out in Vancouver (Canada), found that flowers with large UV patterns received significantly more visits (Figure 4a). Vancouver is outside the natural range of *H. annuus*, suggesting that our results are unlikely to be affected by learned preferences (i.e. pollinators preferring UV patterns they are familiar with in sunflower). While this experiment revealed a clear pattern of pollinator preferences, it involved plants from only two different populations, and effects of other unmeasured factors unrelated to UV pigmentation on visitation patterns cannot be excluded. Therefore, we monitored pollinator visitation in plants grown in a field including 1484 individuals from 106 *H. annuus* populations, spanning the entire range of the species. Within this field, we selected 82 plants, from 49 populations, which flowered at roughly the same time and had comparable numbers of flowers. We divided these plants into three categories, based on the species-wide distribution of LUVp values in *H. annuus* (Figure 1d): small (0-0.3); intermediate (0.5-0.8) and large (>0.95) LUVp. Plants with intermediate UV patterns had the highest visitation rates (Figure 4b; Figure 4 – figure supplement 1). Visitation to plants with small or large UV patterns was less frequent, and particularly low for plants with very small LUVp values (<0.15). A strong reduction in pollination would be expected to result in lower fitness, and to be negatively selected; accordingly, plants with such small LUVp values were rare (∼1.5% of the individuals grown). These results confirm that floral UV patterns play a major role in pollinator attraction, as has been already extensively reported (Horth et al., 2014, Koski and Ashman, 2014, Rae and Vamosi, 2013, Sheehan et al., 2016). They also agree with previous findings in other plant species suggesting that intermediate-to-large UV bullseyes are preferred by pollinators, and only very small UV bullseyes are maladaptive (Horth et al., 2014, Koski and Ashman, 2014). While we cannot exclude that smaller UV bullseyes would be preferred by pollinators in some parts of the *H. annuus* range, this does not seem likely; the most common pollinators of sunflower are ubiquitous across the range of *H. annuus*, and many bee species known to pollinate sunflower are found in both regions where *H. annuus* populations have large LUVp and regions where they have small LUVp (Hurd Jr et al., 1980). While acting as visual cues for pollinators is therefore clearly a major function of floral UV bullseyes, it is unlikely to (fully) explain the patterns of variation that we observe for this trait.

**Figure 4:**
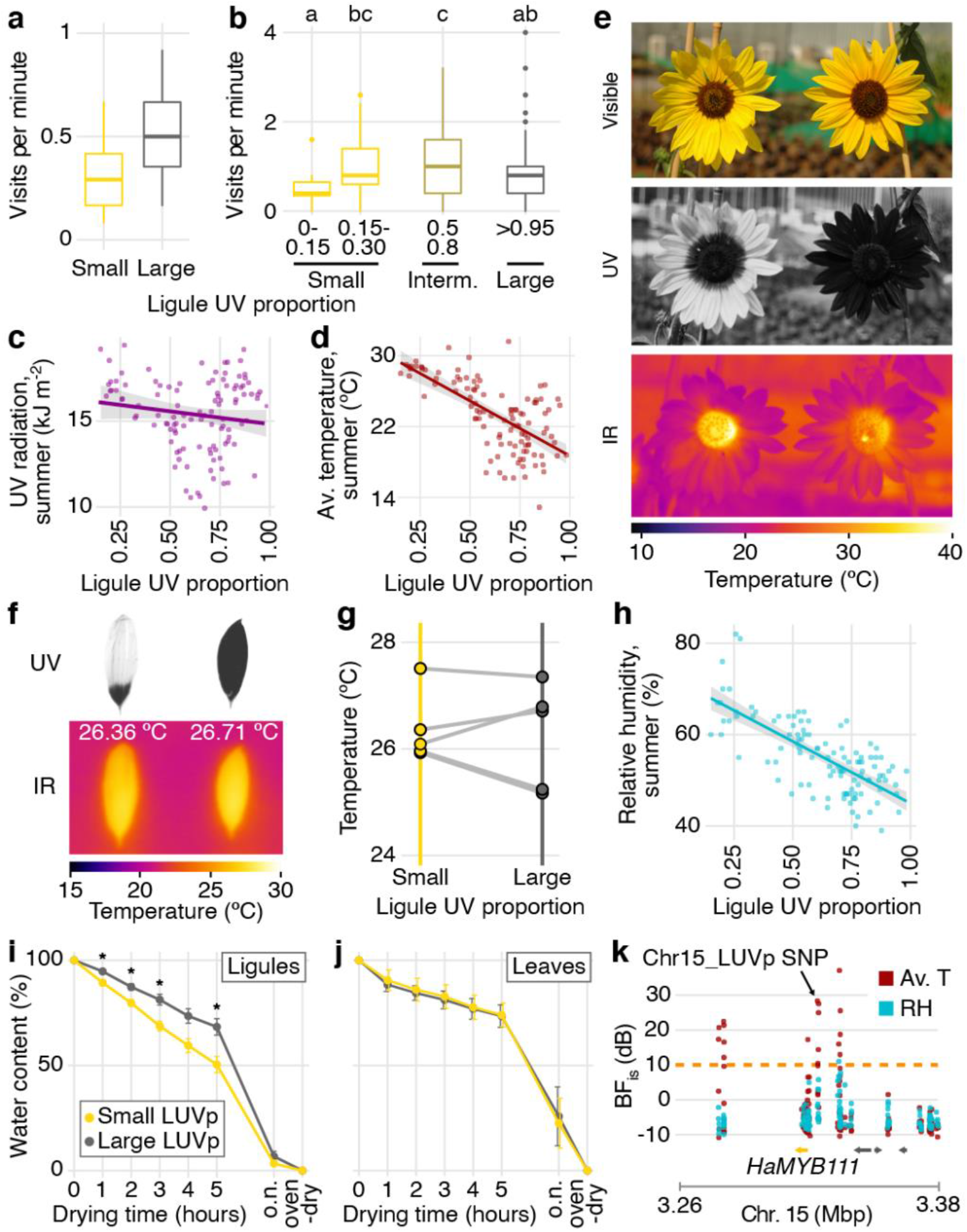
Accumulation of UV pigments in flowers affects pollinator visits and transpiration rates. **a,** Rates of pollinator visitation measured in Vancouver in 2017 (*P* = 0.017; Mann-Whitney U-tests, W = 150, two-sided; *n* = 143 pollinator visits) and **b,** in 2019 (differences between LUVp categories are significant for *P* = 0.0058, Kruskal-Wallis test, *χ*^2^ = 14.54, df = 4; *n* = 1390 pollinator visits. Letters identify groups that are significantly different for *P* < 0.05 in pairwise comparisons, Wilcoxon rank sum test. Exact p-values are reported in the source data for this figure). Boxplots show the median, box edges represent the 25^th^ and 75^th^ percentiles, whiskers represent the maximum/minimum data points within 1.5x interquartile range outside box edges. **c,** Correlation between average LUVp for different populations of *H. annuus* and summer UV radiation (*R^2^* = 0.01, *P* = 0.12) or **d,** summer average temperature (*R^2^* = 0.44, *P* =2.4 × 10^-15^). Grey areas represent 95% confidence intervals. **e,** Sunflower inflorescences pictured in the visible, UV and infrared (IR) range. In the IR picture, a bumblebee is visible in the inflorescence with large LUVp (right). The higher temperature in the centre (disk) of the inflorescence with small LUVp does not depend on ligule UV patterns (Figure 4 – support figure 3). **f,** *H. annuus* ligules after having been exposed to sunlight for 15 minutes. **g,** Pairs of ligules from different sunflower lines were exposed to sunlight for 15 minutes, and their average temperature was measured from IR pictures. **h,** Correlation between average LUVp in *H. annuus* populations and summer relative humidity (*R^2^* = 0.51, *P* = 1.4 × 10^-18^). The grey area represents the 95% confidence interval. **i,**Rate of water loss from ligules and **j,**leaves of wild *H. annuus* plants with large or small LUVp. Values reported are means ± standard error of the mean. *n* = 16 (ligules) or 15 (leaves). Ligules and leaves were left to air-dry and weighed every hour for five hours, after they were left to air-dry overnight (o.n.), and after they were incubated in an oven to remove any residual humidity (oven-dry). Asterisks denote significant differences (p < 0.05, two-sided Welch t-test; exact p-values are reported in the source data for this figure). **k,**GEA for summer average temperature (Av. T) and summer relative humidity (RH) in the *HaMYB111* region. The dashed orange line represents Bayes Factor (BFis) = 10 deciban (dB). GEAs were calculated using two-sided XtX statistics*. n* = 71 populations.

In recent years, the importance of non-pollinator factors in driving selection for floral traits has been increasingly recognized (Strauss and Whittall, 2006). Additionally, flavonol glycosides, the pigments responsible for floral UV patterns in sunflower, are known to be involved in responses to several abiotic stressors (Korn et al., 2008, Nakabayashi et al., 2014b, Pollastri and Tattini, 2011, Schulz et al., 2015). Therefore, we explored whether some of these stressors could drive diversification in floral UV pigmentation. An intuitively strong candidate is UV radiation, which can be harmful to plant cells (Stapleton, 1992). Variation in the size of UV bullseye patterns across the range of *Argentina anserina* (a member of the *Rosaceae* family) has been shown to correlate positively with intensity of UV radiation. Flowers of this species are bowl-shaped, and larger UV-absorbing regions have been proposed to protect pollen from UV damage by absorbing UV radiation that would otherwise be reflected toward the anthers (Koski and Ashman, 2015). However, sunflower inflorescences are much flatter than *A. anserina* flowers, making it unlikely that any significant amount of UV radiation would be reflected from the ligules towards the disk flowers. Studies in another plant with non-bowl-shaped flowers (*Clarkia unguiculata*) have found no evidence of an effect of floral UV patterns in protecting pollen from UV damage (Peach et al., 2020). Consistent with this, the associations between the intensity of UV radiation at our collection sites and floral UV patterns in *H. annuus* was weak (*H. annuus*: *R^2^* = 0.01, *P* = 0.12; Figure 4c; Figure 4 – figure supplement 2).

Across the *Potentillae* tribe (*Rosaceae*), floral UV bullseye size is also weakly associated with UV radiation, but is more strongly correlated with temperature, with lower temperatures being associated with larger UV bullseyes (Koski and Ashman, 2016). We found a similar, strong correlation with temperature in our dataset, with average summer temperatures explaining a large fraction of the variation in LUVp in *H. annuus* (*R^2^* = 0.44, *P* = 2.4 × 10^-15^; Figure 4d; Figure 4 – figure supplement 2). It has been suggested that the radiation absorbed by floral UV pigments could contribute to increasing the temperature of the flower, similar to what has been observed for visible pigments (Koski et al., 2020). This possibility is particularly intriguing for sunflower, in which flower temperature plays an important role in pollinator attraction; inflorescences of cultivated sunflowers consistently face East so that they warm up faster in the morning, making them more attractive to pollinators (Atamian et al., 2016). Larger UV bullseyes could therefore contribute to increasing temperature of the sunflower inflorescences, and their attractiveness of sunflowers to pollinators, in cold climates. However, different levels of UV pigmentation had no effect on the temperature of inflorescences or individual ligules exposed to sunlight (Figure 4e-g; Figure 4 – figure supplement 3). This is perhaps not surprising, given that UV wavelengths represents only a small fraction (3-7%) of the solar radiation reaching the Earth surface (compared to >50% for visible wavelengths), and are therefore unlikely to provide sufficient energy to significantly warm up the ligules (Nunez et al., 1994).

While several geoclimatic variables are correlated across the range of wild *H. annuus*, the single variable explaining the largest proportion of the variation in floral UV patterns in this species was summer relative humidity (RH; *R^2^* = 0.51, *P* = 1.4 × 10^-18^; Figure 4h; Figure 4 – figure supplement 2), with lower humidity being associated with larger LUVp values. Lower relative humidity is generally associated with higher transpiration rates in plants, leading to increased water loss. Flavonol glycosides are known to play an important role in responses to drought stress (Nakabayashi et al., 2014a); in particular, Arabidopsis lines that accumulate higher concentrations of flavonol glycosides due to over-expression of *AtMYB12* lose water and desiccate at slower rates than wild-type plants (Nakabayashi et al., 2014b). Similarly, we found that completely UV-absorbing ligules desiccate significantly slower than largely UV-reflecting ligules (Figure 4i). This difference is not due to general differences in transpiration rates between genotypes, since we observed no comparable trend for rates of leaf desiccation in the same set of sunflower lines (Figure 4j). Transpiration from flowers can be a major source of water loss for plants, and this is known to drive, within species, the evolution of smaller flowers in populations living in dry locations (Galen, 2000, Herrera, 2005, Lambrecht, 2013, Lambrecht and Dawson, 2007). Thus, variation in floral UV pigmentation in sunflowers is likely similarly driven by the role of flavonol glycosides in reducing water loss from ligules, with larger floral UV patterns helping prevent drought stress in drier environments.

One of the main roles of transpiration in plants is facilitating heat dispersion at higher temperatures through evaporative cooling (Burke and Upchurch, 1989, Drake et al., 2018), which could explain the strong correlation between LUVp and temperature across the range of *H. annuus* (Figure 4d). Consistent with this, summer relative humidity and summer temperatures together explain a considerably larger fraction of the variation for LUVp in *H. annuus* than either variable alone (*R^2^* = 0.63; *P* = 0.0017; Figure 1 – source data 1), with smaller floral UV patterns being associated with higher relative humidity and higher temperatures (Figure 4 – figure supplement 2). Despite a more limited range of variation for LUVp, the same trend is present also in *H. petiolaris* (Figure 4 – figure supplement 4). Consistent with a role of floral UV pigmentation in the plant’s response to variation in both humidity and temperature, we found strong associations (dB > 10) between SNPs in the *HaMYB111* region and these variables in genotype-environment association (GEA) analyses (Figure 4k; Figure 4 – source data 3).

### Conclusions

Connecting adaptive variation to its genetic basis is one of the main goals of evolutionary biology. Here, we show that regulatory variation at a single major gene, the transcription factor *HaMYB111*, underlies most of the variation for floral UV patterns in wild *H. annuus*, and that these UV patterns not only have a strong effect on pollinator visits, but they also co-vary with geoclimatic variables (especially relative humidity and temperature) and affect desiccation rates in ligules. By reducing the amount of transpiration in environments with lower relative humidity, UV-absorbing pigments in the ligule help prevent excessive water loss and maintain ligule turgidity; in humid, hot environments (e.g. Southern Texas), lower accumulation of flavonol glycosides would instead promote transpiration from ligules, keeping them cool and avoiding over-heating. The presence of UV pigmentation in the petals of Arabidopsis (also controlled by the Arabidopsis homolog of *MYB111*) further points to a more general protective role of these pigments in flowers, since pollinator attraction is likely not critical for fertilization in this largely selfing species. Additionally, a role in reducing water loss from petals is consistent with the overall trend in increased size of floral UV patterns over the past 80 years that has been observed in herbarium specimens (Koski et al., 2020); due to changing climates, relative humidity over land has been decreasing in recent decades, which could result in higher transpiration rates (Byrne and O’Gorman, 2018). Further studies will be required to confirm the existence of this trend and assess its strength.

More generally, our study highlights the complex nature of adaptive variation, with selection pressures from both biotic and abiotic factors shaping the patterns of diversity that we observe across natural populations. Floral diversity in particular has long been attributed to the actions of animal pollinators. Our work adds to a growing literature demonstrating the contributions of abiotic factors, most notably drought and heat stress, to this diversity. In sum, it is not all about sex, even for flowers.

## Methods

### Plant material and growth conditions

Sunflower lines used in this paper were grown from seeds collected from wild populations (Todesco et al., 2020), or obtained from the North Central Regional Plant Introduction Station in Ames, Iowa, USA. Sunflower seeds were surface sterilized by immersion for 10 minutes in a 1.5% sodium hypochlorite solution. Seeds were then rinsed twice in distilled water and treated for at least one hour in a solution of 1% PPM (Plant Cell Technologies, Washington, DC, USA), a broad-spectrum biocide/fungicide, to minimize contamination, and 0.05 mM gibberellic acid (Sigma-Aldrich, St. Louis, MO, USA). They were then scarified, de-hulled, and kept for two weeks at 4 °C in the dark on filter paper moistened with a 1% PPM solution. Following this, seeds were kept in the dark at room temperature until they germinated. For common garden experiments, the seedlings were then transplanted in peat pots, grown in a greenhouse for two weeks, then moved to an open-sided greenhouse for a week for acclimation, and finally transplanted in the field. For all other experiments seedlings were transplanted in 2-gallon pots filled with Sunshine #1 growing mix (Sun Gro Horticulture Canada, Abbotsford, BC, Canada). For the wild sunflower species shown in Figure 1 – figure supplement 1b, following sterilization, seeds were scarified and then dipped in fuscicosin solution (1.45 µM) for 15 minutes, dehulled, germinated in the dark for at least 8-10 days, and then grown in pots for three weeks before transplanting into 2-gallon pots filled with a blend of sandy loam, organic compost and mulch. Those plants were grown at the UC Davis field experiment station (California, USA) from July to October 2017. A complete list of sunflower accessions and their populations of origin is reported in Figure 1 - source data 1 and Figure 1 - source data 2.

Seeds from the following Arabidopsis lines were obtained from the Arabidopsis Biological Resource Center: Col-0 (CS28167), *myb111* (CS9813), *myb12* (CS9602) and *myb12/myb111* (CS9980). Seeds were stratified in 0.1% agar at 4 °C in the dark for four days, and then sown in pots containing Sunshine #1 growing mix. Plants were grown in growth chambers at 23 °C in long-day conditions (16 h light, 8 h dark).

### Common garden

Two common garden experiments were performed, in 2016 and 2019. After germination and acclimation, plants were transplanted at the Totem Plant Science Field Station of the University of British Columbia (Vancouver, Canada). In the 2016 common garden experiment, each sunflower species was grown in a separate field. Pairs of plants from the same population were randomly distributed within each field. In the 2019 common garden experiment, plants were sown using a completely randomized design.

In the summer of 2016, ten plants from each of 151 selected populations of wild *H. annuus*, *H. petiolaris*, *H. argophyllus* and *H. niveus* were grown. Plants were transplanted in the field on the 25^th^ of May (*H. argophyllus*), 2^nd^ of June (*H. petiolaris* and *H. niveus*) and 7^th^ of June 2016 (*H. annuus*). Up to four inflorescences from each plant were collected for visible and UV photography. In the summer of 2019, fourteen plants from each of 106 populations of wild *H. annuus* were transplanted in the field on 6^th^ of June. These included 65 of the populations grown in the previous common garden experiment, and 41 additional populations that were selected to complement their geographic distribution. At least two ligules from different inflorescences for each plant were collected for visible and UV photography.

Sample size for the common garden experiments was determined by the available growing space and resources. Ten-to-fourteen individuals were grown for each population because this would provide a good representation of the variation present in each population, while maximizing the number of populations that could be surveyed.

Researcher were not blinded as to the identity of individual samples. However, information about their populations of origin and/or LUVp phenotypes were not attached to the samples during data acquisition.

### Ultraviolet and infrared photography

Ultraviolet patterns were imaged in whole flowerheads or detached ligules using a Nikon D70s digital camera, fitted with a Noflexar 35-mm lens and a reverse-mounted 2-inch Baader U-Filter (Baader Planetarium, Mammendorf, Germany), which only allows the transmission of light between 320 and 380 nm. Wild sunflower species shown in Figure 1 – figure supplement 1b were imaged using a Canon DSLR camera in which the internal hot mirror filter had been replaced with a UV bandpass filter (LifePixel, Mukilteo, WA). The length of the whole ligule (L_L_) and the length of the UV-absorbing part at the base of the ligule (L_UV-abs_) were measured using Fiji (Schindelin et al., 2012, Schneider et al., 2012). Ligule UV proportion was measured as the ratio between the two (LUVp = L_UV-abs_/L_L_). In some *H. annuus* individuals, the upper, “UV-reflecting” portion of the ligules (L_UV-ref_) also displayed a lower level of UV-absorption; in those cases, these regions were weighted at 50% of fully UV-absorbing regions, using the formula LUVp = (L_UV-abs_/L_L_) + ½(L_UV-ref_/L_L_). For pictures of whole inflorescences, LUVp values were averaged from three different ligules. For each individual, LUVp values were averaged between all the inflorescences or detached ligules measurements.

Infrared pictures were taken using a Fluke TiX560 thermal imager (Fluke Corporation, Everett, WA, USA) and analyzed using the Fluke Connect software (v1.1.536.0). For time series experiments, plants were germinated as above (see “Common Garden”), grown in 2-gallon pots in a greenhouse until they produced four true leaves, and then moved to the field. On three separate days in August 2017, pairs of inflorescences with opposite floral UV patterns at similar developmental stages were selected and made to face East. Infrared images were taken just before sunrise, ∼5 minutes after sunrise, and then at 0.5, 1, 2, 3 and 4 hours after sunrise. For infrared pictures of detached ligules, plants were grown in a greenhouse. Flowerheads were collected and kept overnight in a room with constant temperature of 21 °C, with their stems immersed in a beaker containing distilled water. The following day, pairs of ligules with contrasting LUVp were removed and arranged on a sheet of white paper. Infrared pictures were taken immediately before exposing the ligules to the sun, and again 15 minutes after that.

### Library preparation, sequencing and SNP calling

Whole-genome shotgun (WGS) sequencing library preparation and sequencing, as well as SNP calling and variant filtering, for the *H. annuus* and *H. petiolaris* individuals used for GWA analyses in this paper were previously described (Todesco et al., 2020). Briefly, DNA was extracted from leaf tissue using a modified CTAB protocol (Murray and Thompson, 1980, Zeng et al., 2002), the DNeasy Plant Mini Kit or a DNeasy 96 Plant Kit (Qiagen, Hilden, Germany). Genomic DNA was sheared to an average fragment size of 400 bp using a Covaris M220 ultrasonicator (Covaris, Woburn, MA, USA). Libraries were prepared using a protocol largely based on (Rowan et al., 2015), the TruSeq DNA Sample Preparation Guide from Illumina (Illumina, San Diego, CA, USA) and (Rohland and Reich, 2012), with the addition of an enzymatic repeats depletion step using a Duplex-Specific Nuclease (DSN; Evrogen, Moscow, Russia) (Matvienko et al., 2013, Shagina et al., 2010, Todesco et al., 2020). All libraries were sequenced at the McGill University and Génome Québec Innovation Center on HiSeq2500, HiSeq4000 and HiSeqX instruments (Illumina) to produce paired end, 150 bp reads.

Sequences were trimmed for low quality using Trimmomatic (v0.36) (Bolger et al., 2014) and aligned to the *H. annuus* XRQv1 genome (Badouin et al., 2017) using NextGenMap (v0.5.3) (Sedlazeck et al., 2013). We followed the best practices recommendations of The Genome Analysis ToolKit (GATK) (Poplin et al., 2017) and executed steps documented in GATK’s germline short variant discovery pipeline (for GATK 4.0.1.2). During genotyping, to reduce computational time and improve variant quality, genomic regions containing transposable elements were excluded (Badouin et al., 2017). We then used GATK’s VariantRecalibrator (v4.0.1.2) to select high quality variants. SNP data were then filtered for minor allele frequency (MAF) ≥ 0.01, genotype rate ≥ 90%, and to keep only bi-allelic SNPs.

Filtered SNPs were then re-mapped to the improved reference assembly HA412-HOv2 (Staton and Lázaro-Guevara, 2020) using BWA (v0.7.17) (Li, 2013). These re-mapped SNPs were used for all analyses, excluding the GWA for the region surrounding the *HaMYB111* locus that used un-filtered variants based on the XRQv1 assembly (Figure 2 – figure supplement 3).

The SNP dataset used to determine the genotype at the Chr15_LUVp SNP in other species (*H. argophyllus*, *H. niveus*, *H. debilis*, *H. decapetalus*, *H. divaricatus* and *H. grosseserratus*) was based on WGS data generated for (Todesco et al., 2020) and is described in (Owens et al., 2021). Sequence data for the Sunflower Association Mapping population was reported in (Hubner et al., 2019).

### Genome-wide association mapping

Genome-wide association analyses for LUVp were performed for *H. annuus, H. petiolaris petiolaris* and *H. petiolaris fallax,* using two-sided mixed models implemented in EMMAX (v07Mar2010) (Kang et al., 2010) or in the EMMAX module in EasyGWAS (Grimm et al., 2017). For all runs, the first three principal components (PCs) were included as covariates, as well as a kinship matrix. Only SNPs with MAF >5% were included in the analyses. Sample size was estimated to be sufficient to provide an 85% probability of detecting loci explaining 5% or more of the phenotypic variance in *H. annuus*, 8% of variance in *H. petiolaris*.

### F_2_ populations and genotyping

Individuals from population ANN_03 from California, USA (large LUVp) and ANN_55 from Texas, USA (small LUVp) were grown in 2-gallon pots in a field. When the plants reached maturity, they were moved to a greenhouse, where several inflorescences were bagged and crossed. The resulting F_1_ seeds were germinated and grown in a greenhouse, and pairs of siblings were crossed (wild sunflowers are overwhelmingly self-incompatible). The resulting F_2_ populations were grown both in a greenhouse in the winter of 2019 (*n* = 42 individuals for population 1, 38 individuals for population 2) and in a field as part of the 2019 common garden experiments (*n* = 54 individuals for population 1, 50 individuals for population 2). DNA was extracted from young leaf tissue as described above. All plants were genotyped for the Chr15_LUVp SNPs using a custom TaqMan SNP genotyping assay (Thermo Fisher Scientific, Walthman, MA, USA) on a Viia 7 Real-Time PCR system (Thermo Fisher Scientific).

### Metabolite analyses

Methanolic extractions were performed following (Stracke et al., 2007). Sunflower ligules (or portions of them) and Arabidopsis petals were collected and flash-frozen in liquid nitrogen. The tissue was ground to a fine powder by adding 10-15 zirconia beads (1 mm diameter) and using a TissueLyser (Qiagen) for sunflower ligules, or using a plastic pestle in a 1.5 ml tube for Arabidopsis petals. 0.5 ml of 80% methanol were added, and the samples were further homogenized and incubated at 70 °C for 15 minutes. They were then centrifuged at 15.000g for 10 minutes, and the supernatant was dried in a SpeedVac (Thermo Fisher Scientifics) at 60 °C. Samples were then resuspended in 1 µl (sunflower) or 2.5 µl (Arabidopsis) of 80% methanol for every mg of starting tissue.

The extracts were analyzed by LC/MS/MS using an Agilent 1290 UHPLC system (Agilent Technologies, Santa Clara, CA, USA) coupled with an Agilent 6530 Quadrupole Time of Flight mass spectrometer. The chromatographic separation was performed on Atlantis T3-C18 reversed-phase (50 mm × 2.1 mm, 3 µm) analytical columns (Waters Corp, Milford, MA, USA). The column temperature was set at 40 °C. The elution gradient consisted of mobile phase A (water and 0.2% formic acid) and mobile phase B (acetonitrile and 0.2% formic acid). The gradient program was started with 3% B, increased to 25% B in 10 min, then increased to 40% B in 13 min, increased to 90% B in 17 min, held for 1 min and equilibrated back to 3% B in 20 min. The flow rate was set at 0.4 mL/min and injection volume was 1 µl. A PDA (photo diode array) detector was used for detection of UV-absorption in the range of 190-600 nm.

MS and MS/MS detection were performed using an Agilent 6530 accurate mass Quadrupole Time of Flight mass spectrometer equipped with an ESI (electrospray) source operating in both positive and negative ionization modes. Accurate positive ESI LC/MS and LC/MS/MS data were processed using the Agilent MassHunter software to identify the analytes. The ESI conditions were as follows: nebulizing gas (nitrogen) pressure and temperature were 30 psi and 325 °C; sheath gas (nitrogen) flow and temperature were 12 L/min, 325 °C; dry gas (nitrogen) was 7 l/min. Full scan mass range was 50-1700 m/z. Stepwise fragmentation analysis (MS/MS) was carried out with different collision energies depending on the compound class.

### Transgenes and expression assays

Total RNA was isolated from mature and developing ligules, or part of ligules, using TRIzol (Thermo Fisher Scientific) and cDNA was synthesized using the RevertAid First Strand cDNA Synthesis kit (Thermo Fisher Scientific). Genomic DNA was extracted from leaves of Arabidopsis using CTAB (Murray and Thompson, 1980). A 1959 bp-long fragment (*pAtMYB111*) from the promoter region of *AtMYB111* (*At5g49330*), including the 5’-UTR of the gene, was amplified using Phusion High-Fidelity DNA polymerase (New England Biolabs, Ipswich, MA, USA) and introduced in pFK206 derived from pGREEN (Hellens et al., 2000). Alleles of *HaMYB111* were amplified from cDNA from ligules, and placed under the control of *pAtMYB111* (in the plasmid describe above) or of the constitutive *CaMV 35S* promoter (in pFK210, derived as well from pGREEN (Hellens et al., 2000)). Constructs were introduced into *A. thaliana* plants by *Agrobacterium tumefaciens*-mediated transformation (strain GV3101) (Weigel and Glazebrook, 2002). At least five independent transgenic lines with levels of UV pigmentation comparable to the ones shown in Figure 3d were recovered for each construct. For expression analyses, qRT-PCRs were performed on cDNA from ligules using the SsoFast EvaGreen Supermix (Bio-Rad, Hercules, CA, USA) on a CFX96 Real-Time PCR Detection System (Bio-Rad). Expression levels were normalized against *HaEF1α*. For the expression analyses shown in Figure 3h,i, portions of ligules were collected at different developmental stages from three separate inflorescences from one individuals for each species (biological replicates). Three qRT-PCRs were run for each sample (technical replicates). For the expression analysis shown in Fig. 3j, samples were collected from wild *H. annuus* individuals grown as part of the 2019 common garden experiment. Ligules were collected from developing inflorescences of 24 individuals with large LUVp (from 10 populations) and 22 individuals with small LUVp (from 11 populations). Three technical replicates were performed for each individual. Sample size for this experiment was determined by the number of available plants with opposite LUVp phenotypes and at the appropriate developmental stage. Primers used for cloning and qRT-PCR are given in Table 1.

**Table 1:**
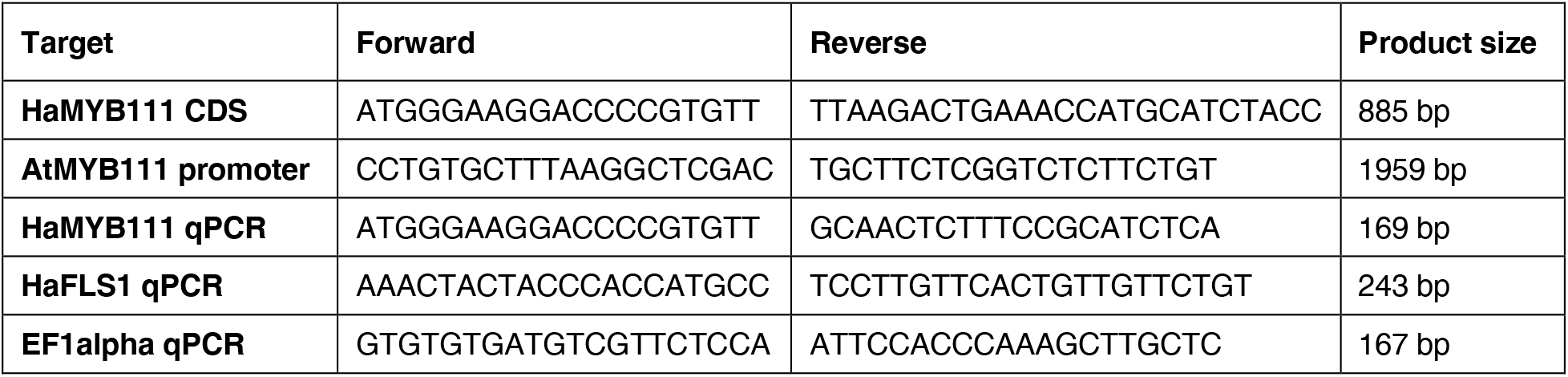
Oligonucleotides used in this study.

### Sanger and PacBio sequencing

Fragments ranging in size from 1.5 to 5.5 kbp were amplified from genomic DNA of 20 individuals that had been previously re-sequenced (Todesco et al., 2020), and whose genotype at the Chr15_LUVp SNP was therefore known, using Phusion High-Fidelity DNA polymerase (New England Biolabs). Fragments were then cloned in either pBluescript or pJET (Thermo Fisher Scientific) and sequenced on a 3730S DNA analyzer using BigDye Terminator v3.1 sequencing chemistry (Applied Biosystem, Foster City, CA, USA).

For long reads sequencing, seed from wild *H. annuus* populations known to be homozygous for different alleles at the Chr15_LUVp SNP were germinated and grown in a greenhouse. After confirming that they had the expected LUVp phenotype, branches from each plant were covered with dark cloth for several days, and young, etiolated leaves were collected and immediately frozen in liquid nitrogen. High molecular weight (HMW) DNA was extracted using a modified CTAB protocol (Stoffel et al., 2012). All individuals were genotyped for the Chr15_LUVp SNP using a custom TaqMan SNP genotyping assay (Thermo Fisher Scientific, see above) on a CFX96 Real-Time PCR Detection System (Bio-Rad). Two individuals, one with large and one with small LUVp, were selected. HiFi library preparation and sequencing were performed at the McGill University and Génome Québec Innovation Center on a Sequel II instrument (PacBio, Menlo Park, CA, USA). Each individual was sequenced on an individual SMRT cell 8M, resulting in average genome-wide sequencing coverage of 6-8X.

### Pollinator preferences assays

In September 2017, pollinator visits were recorded in individual inflorescences of pairs of plants with large and small LUVp grown in pots in a field adjacent the Nursery South Campus greenhouses of the University of British Columbia. Pairs of inflorescences were filmed using a Bushnell Trophy Cam HD (Bushnell, Overland Park, KS, USA) in 12-minute intervals. Visitation rates were averaged over 14 such movies (Figure 4 – source data 1).

In summer 2019, pollinator visits were scored in a common garden experiment consisting of 1484 *H. annuus* plants at the Totem Plant Science Field Station of the University of British Columbia. Over five days, between the 29^th^ of July and the 7^th^ of August, pollinator visits on individual plants were counted over five-minute intervals, for a total of 435 series of measurements on 111 plants from 51 different populations (Figure 4 – source data 1). To generate a more homogenous and comparable dataset, measurements for plants with too few (1) or too many (>10) flowers were excluded from the final analysis.

### Correlations with environmental variables and genotype-environment association analyses

Twenty topo-climatic factors were extracted from climate data collected over a 30-year period (1961-1990) for the geographic coordinates of the population collection sites, using the software package Climate NA (Wang et al., 2016) (Figure 1 – source data 1). Additionally, UV radiation data were extracted from the glUV dataset (Beckmann et al., 2014) using the R package “raster” (Hijmans, 2020, R Core Team, 2020). Correlations between individual environmental variables and LUVp were calculated using the “lm” function implemented in R; a correlation matrix between all environmental variables and LUVp was calculated using the “cor” function in R and plotted using the “heatmap.2” function in the “gplots” package (Warnes et al., 2009).

GEAs were analyzed using BayPass (Gautier, 2015) version 2.1. Population structure was estimated by choosing 10,000 putatively neutral random SNPs under the BayPass core model. The Bayes factor (denoted BF_is_ as in (Gautier, 2015)) was then calculated under the standard covariate mode. For each SNP, BF_is_ was expressed in deciban units [dB, 10 log_10_ (BF_is_)]. Significance was determined following (Gautier, 2015), and employing Jeffreys’ rule (Jeffreys, 1961), quantifying the strength of associations between SNPs and variables as “strong” (10 dB ≤ BF_is_ < 15 dB), “very strong” (15 dB ≤ BF_is_ < 20 dB) and decisive (BF_is_ ≥ 20 dB).

### Desiccation assays

Inflorescence and leaves were collected from well-watered, greenhouse-grown plants, and brought to an environment kept at 21°C. They were left overnight with their stems or petioles immersed in a beaker containing distilled water. The following morning 1-2 leaves from each plant, and three ligules from each inflorescence were individually weighed and hanged to air dry at room temperature (21 °C). Their weight was measured at one-hour intervals for five hours, and then again the following morning. Leaves and ligules were then incubated for 48 hours at 65 °C in an oven to determine their dry weight. Total water content was measured as the difference between the initial fresh weight (W_0_) and dry weight (W_d_). Water loss was expressed as a fraction of the total water content of each organ, using the formula [(W_i_-W_d_)/(W_0_-W_d_)] × 100, where W_i_ is the weight of a sample at a time i. The assay was performed on samples from 16 (ligules) or 15 (leaves) individuals belonging to 7 (ligules) or 8 (leaves) different populations of *H. annuus*.

### Data availability

All raw sequenced data are stored in the Sequence Read Archive (SRA) under BioProjects PRJNA532579, PRJNA398560 and PRJNA736734. Filtered SNP datasets are available at https://rieseberglab.github.io/ubc-sunflower-genome/. Raw sequencing data and SNP datasets have been previously described in (Todesco et al., 2020). The sequences of individual alleles at the *HaMYB111* locus and of *HaMYB111* coding sequences have been deposited at GenBank under accession numbers XXX-XXX and MZA410295-MZA410296, respectively. All other data are available in the main text or in the source data provided with the manuscript.

## Acknowledgments

This research was conducted in the ancestral and unceded territory of the xʷməθkʷəy̓əm (Musqueam) People. We thank Andrea Todesco, Daniela Rodeghiero, Emma Borger, Quinn Anderson, Jennifer Lipka, Jasmine Lai, Hafsa Ahmed, Dominique Skonieczny, Ana Parra, Cassandra Konecny, Kelsie Morioka and Daniel Yang for assistance with field work and data acquisition, Melina Byron and Glen Healy at UBC and the UC Davis Plant Sciences Field Station personnel for assistance with greenhouse and field experiments, Elizabeth Elle and Tyler Kelly for help planning the pollinator preferences experiments, Laura Marek and the USDA-ARS in Ames, IA, USA, for providing sunflower seeds, and Chase Mason for providing cuttings for *Phoebantus tenuifolius*. Maps were realized using tiles from Stamen Design (https://stamen.com), under CC BY 3.0, from data by OpenStreetMaps contributors (https://openstreetmap.org), under ODbL. Funding was provided by Genome Canada and Genome BC (LSARP2014-223SUN), the NSF Plant Genome Program (DBI-1444522 and DBI-1817628), the University of California, Berkeley, and an HFSP long-term postdoctoral fellowship to M.T. (LT000780/2013).

## Competing interests

The authors declare no competing interests.

## Figure supplements

**Figure 1 – figure supplement 1:**
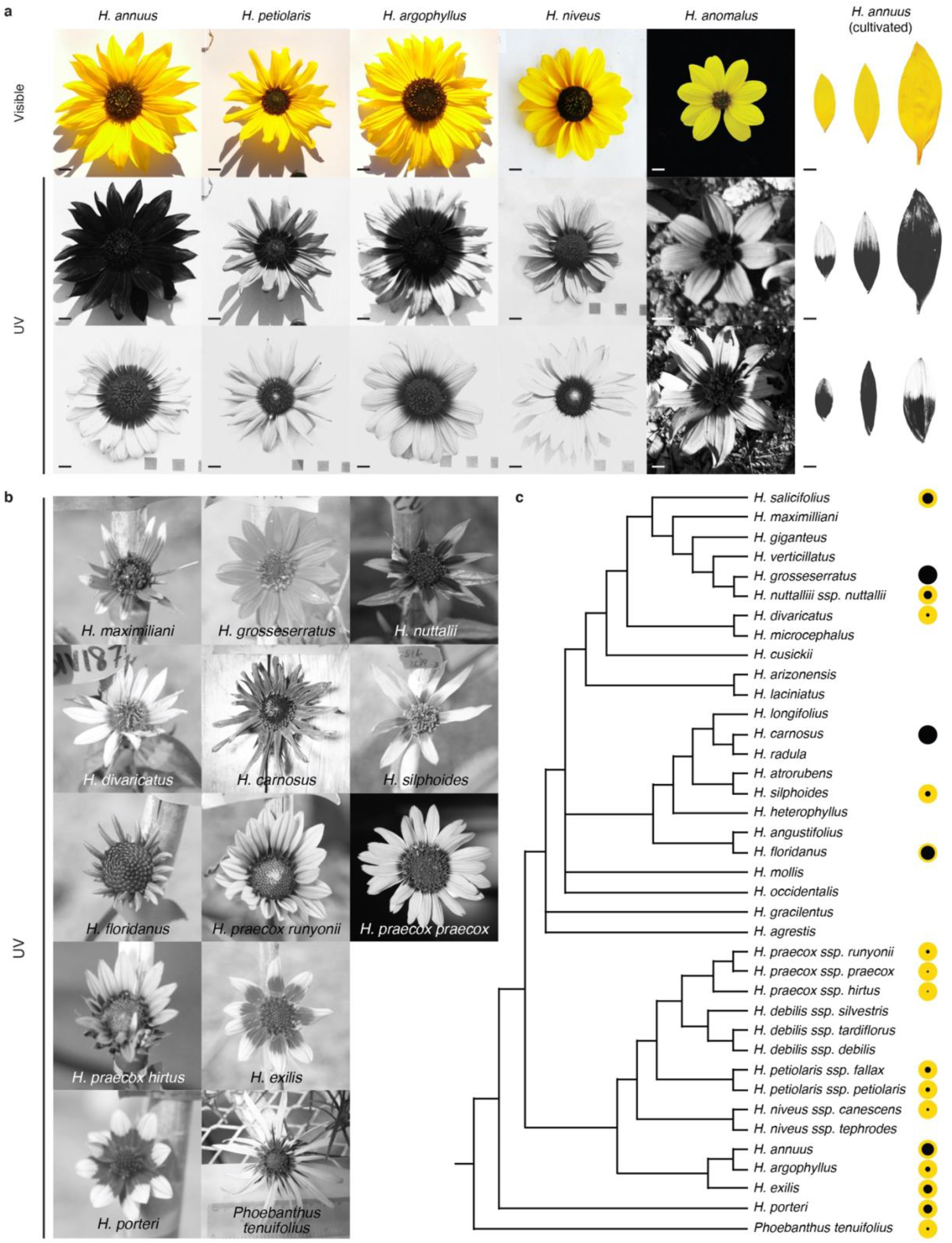
Floral UV patterns in wild sunflower species and cultivated sunflower. **a,** Visible and UV images of inflorescences from five wild sunflower species, and ligules of six cultivated sunflower lines. Variation in floral UV patterns was found within all these species. Scale bar = 1 cm. **b,** UV images of inflorescences from twelve wild sunflower species and for the outgroup *Phoebanthus tenuifolius*. Images are not to scale. **c,**Species tree for 31 sunflower species and *P. tenuifolius*, modified from (Stephens et al., 2015). The size of the black dots to the right of each species name is proportional to the average size of bullseye patterns measured for that species or subspecies (Figure 1 – source data 1). For the species in **a**, bullseye values are averages for ≥42 individuals (see also Figure 1 – figure supplement 2). For the species in **b**, bullseye values are for single individuals or averages for up to three individuals. Two taxa in the original species tree, *H. petiolaris* and *H. neglectus*, were renamed to *H. petiolaris ssp. petiolaris* and *H. petiolaris ssp. fallax*to reflect the current understanding of their identities.

**Figure 1 – figure supplement 2:**
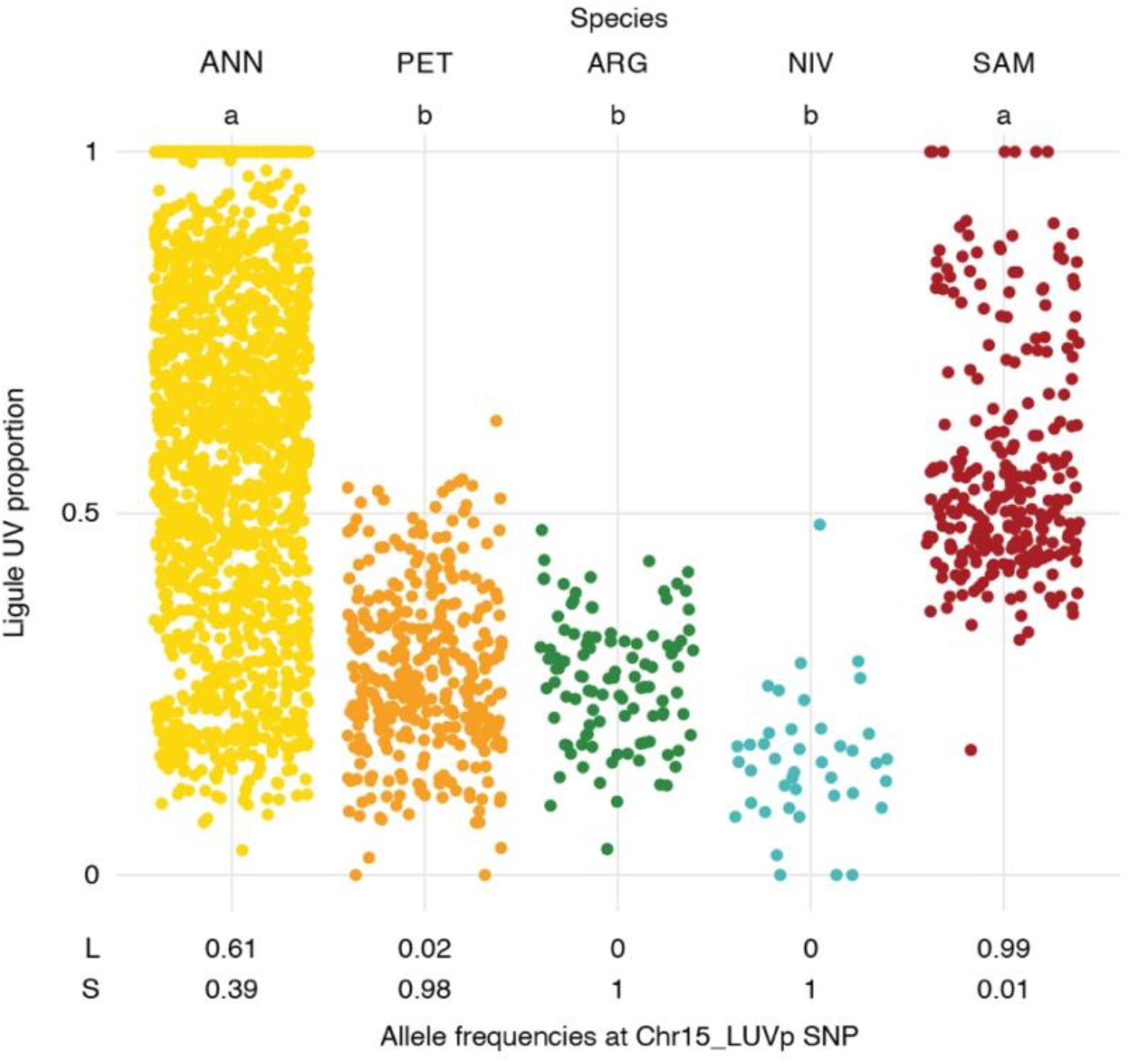
LUVp variation in wild sunflower species and cultivated sunflower. LUVp values for individuals of four wild sunflower species and of the cultivated sunflower association mapping (SAM) population, and allele frequencies at the Chr15_LUVp SNP. ANN = wild *H. annuus* (*n* = 1589 from 110 populations), PET = *H. petiolaris* (*n* = 351 individuals from 40 populations), ARG = *H. argophyllus* (*n* = 105 individuals from 27 populations), NIV = *H. niveus* (*n* = 42 individuals from 9 populations), SAM = cultivated *H. annuus* (*n* = 275 individuals). Letters identify groups that are significantly different for p < 0.001 (one-way ANOVA with post-hoc Tukey HSD test, F = 247, df = 4). Exact p-values for pairwise comparisons are reported in Figure 1 – source data 2.

**Figure 2 – figure supplement 1:**
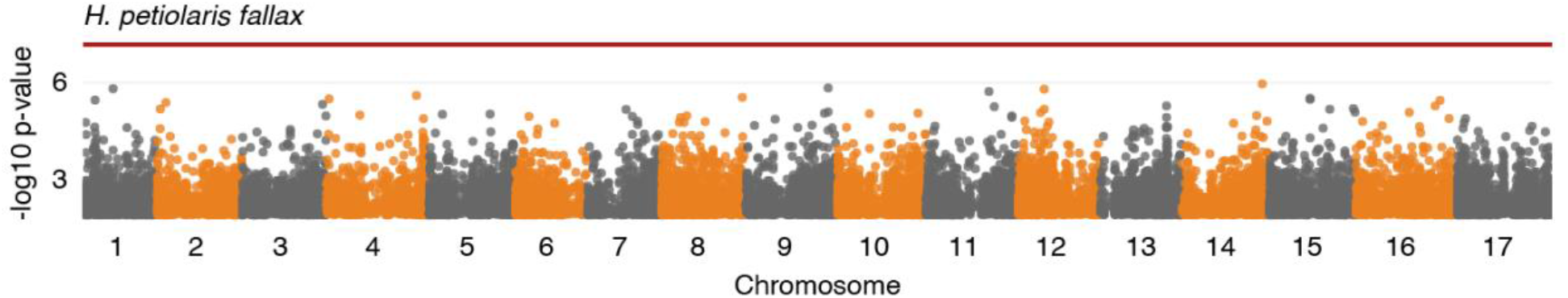
LUVp GWAS in *H. petiolaris fallax.* LUVp GWAS in *H. petiolaris fallax* (*n* = 193 individuals). The red line represents 5% Bonferroni-corrected significance. GWAs were calculated using two-sided mixed models. Only positions with -log10 p-value > 2 are plotted.

**Figure 2 – figure supplement 2:**
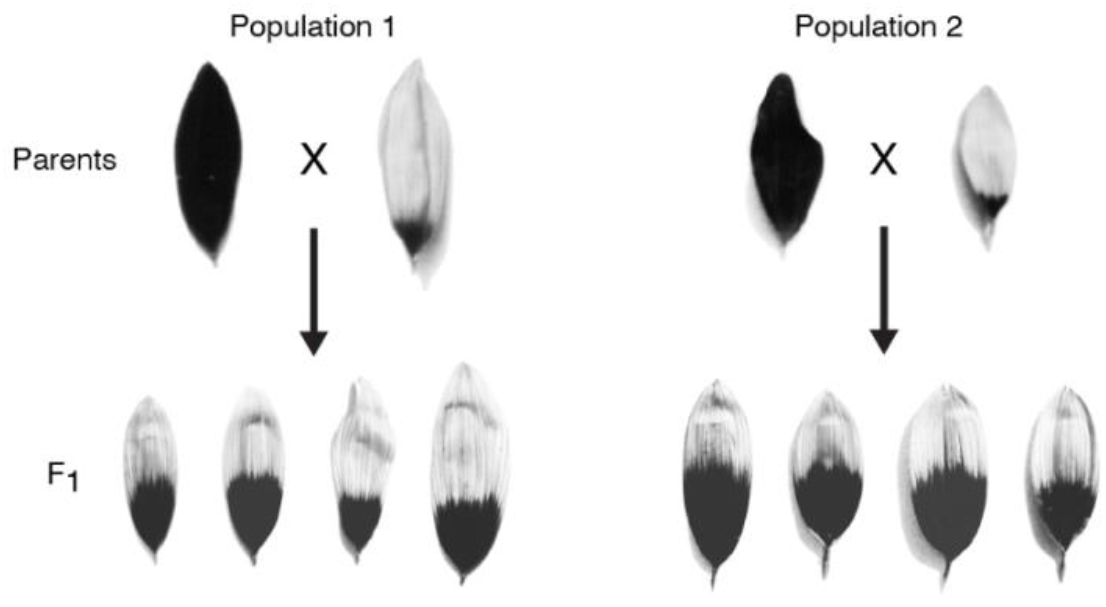
Floral UV patterns in the parental lines of F_2_ populations. UV images of ligules of the parental lines for the F_2_ populations shown in Figure 2e, and their F_1_ progeny. A pair of F_1_ plants was selected and crossed for each population to generate the F_2_ progeny.

**Figure 2 – figure supplement 3:**
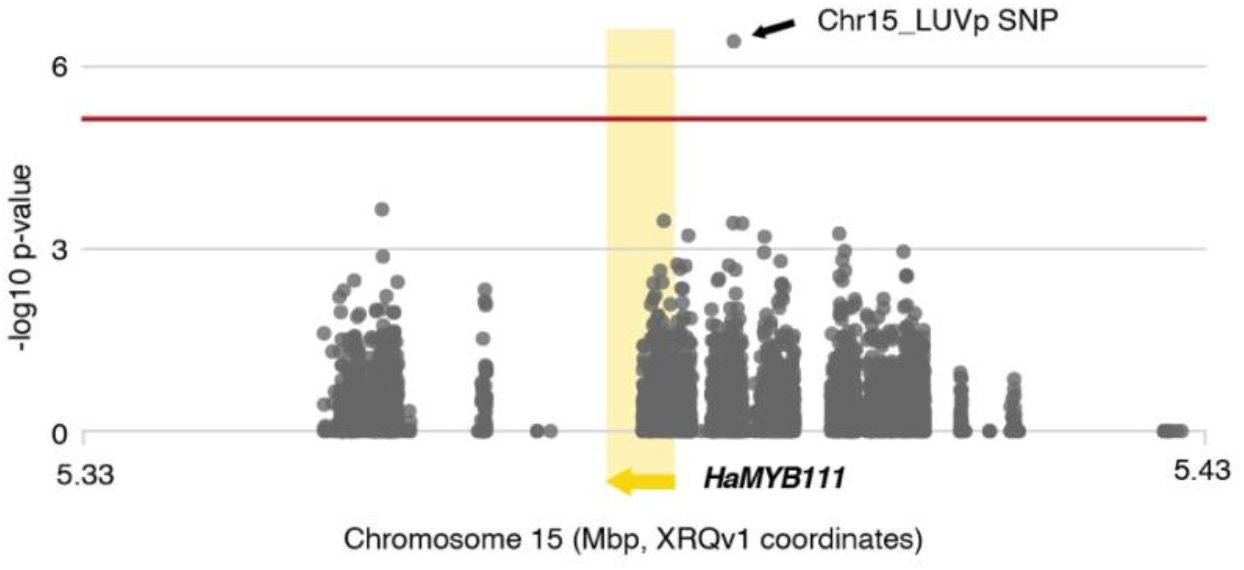
LUVp GWAS in unfiltered *H. annuus* datasets. LUVp GWAS *H. annuus* using an un-filtered variants dataset in a 100 kbp region surrounding *HaMYB111* (*n* = 563 individuals). Relaxing variant filtering parameters, to capture more of the polymorphisms at the *HaMYB111* locus, resulted in an almost 50-fold increase in the number of variants in this region, from 142 to 6949. Regions in which no SNPs are reported contain highly repetitive sequences, and were masked before reads mapping. As re-mapping to the improved HA412v2 reference assembly of the complete *H annuus* set of >222M unfiltered variants would have been computationally intensive, positions are shown based on the original XRQv1 reference assembly. The red line represents 5% Bonferroni-corrected significance. GWAs were calculated using two-sided mixed models.

**Figure 3 – figure supplement 1:**
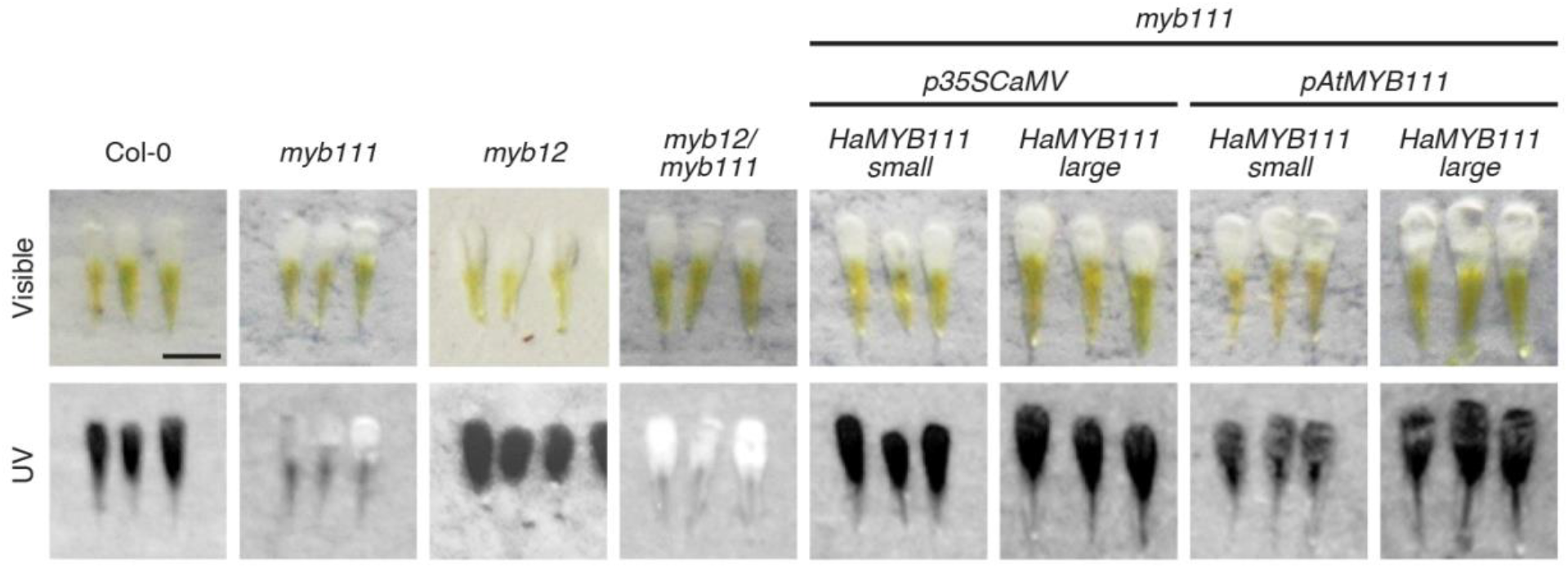
Floral UV patterns in Arabidopsis lines. Visible and UV images of Arabidopsis petals. Col-0 = wild type Arabidopsis. Scale bar = 1 mm.

**Figure 3 – figure supplement 2:**
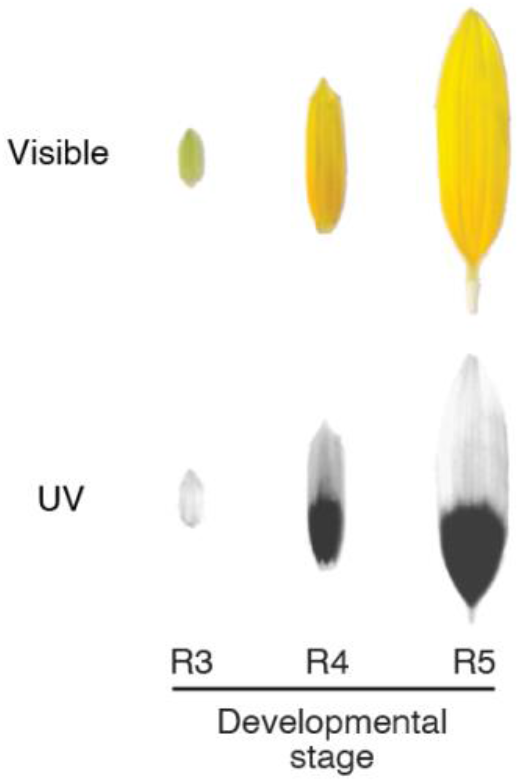
Stages of ligule development in *H. petiolaris*.

**Figure 4 – figure supplement 1:**
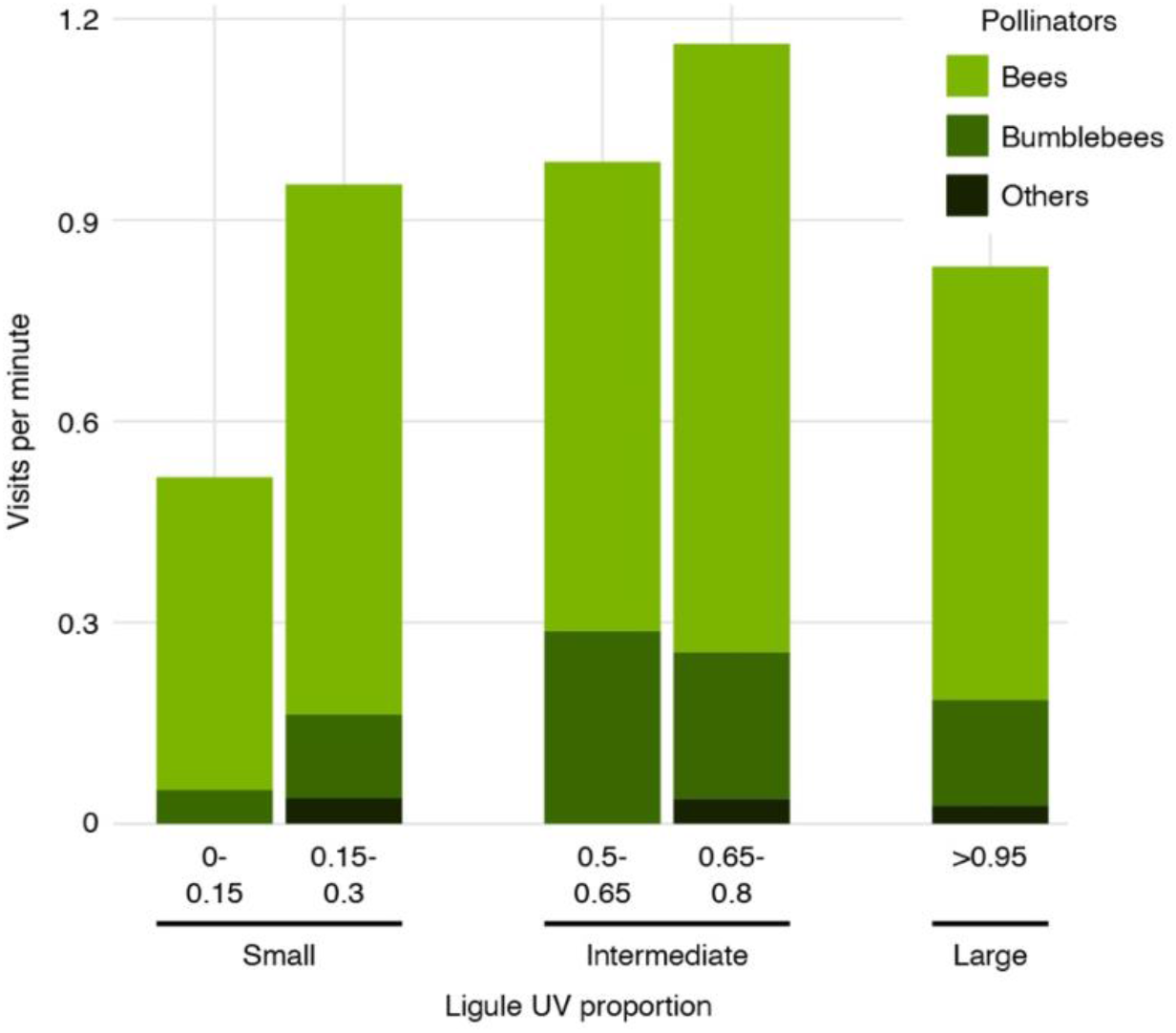
Mean pollinator visits in the 2019 field experiment divided by category of pollinators. “Bees” were exclusively honey bees; “Bumblebees” included several *Bombus* species; “Others” were mostly *Megachile* bees (*n* = 1390 pollinator visits). Pollinators were overwhelmingly bumblebees in the 2017 field experiment.

**Figure 4 – figure supplement 2:**
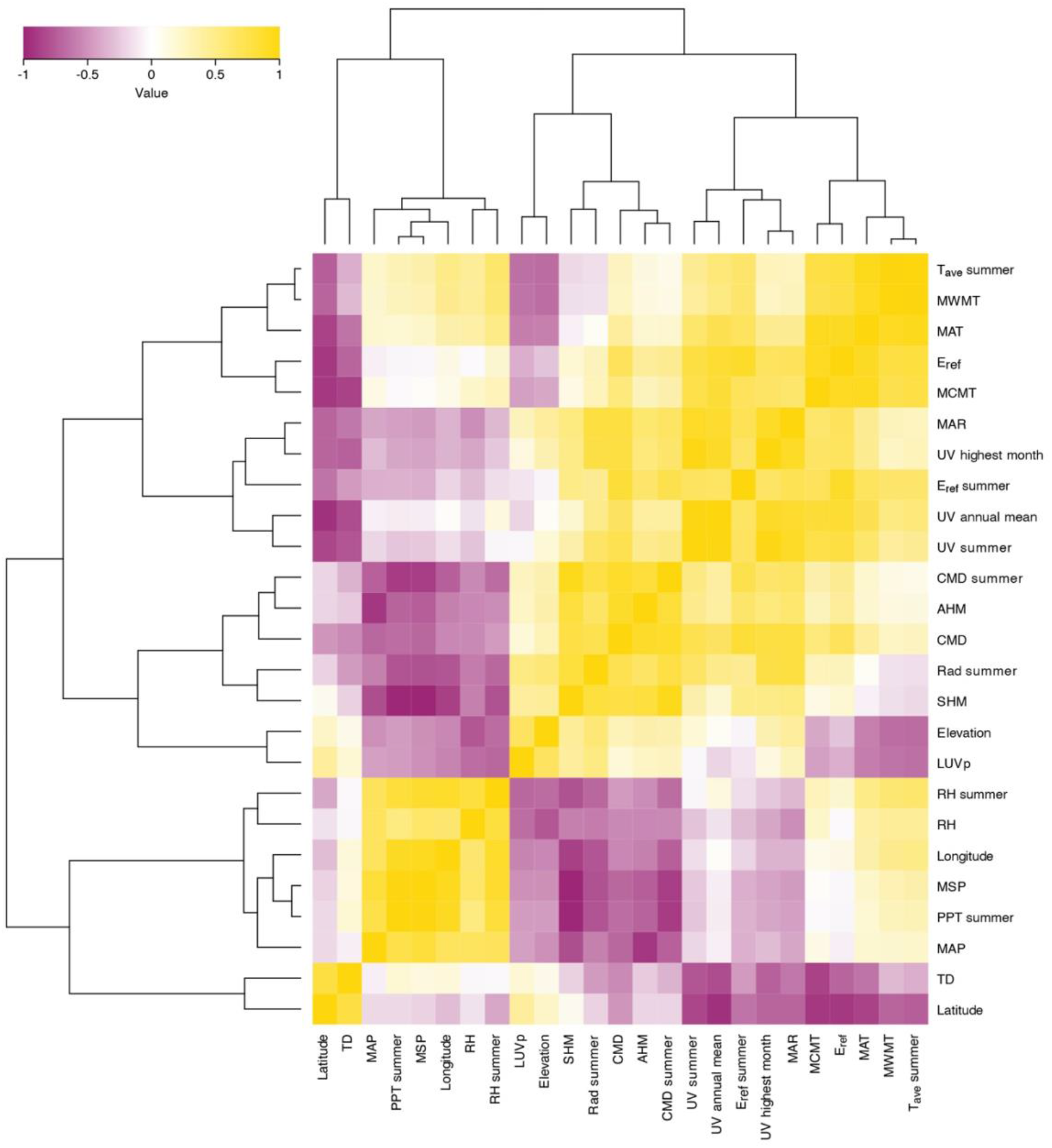
Spearman correlation heatmap for LUVp and environmental variables in wild *H. annuus* populations. Source data for this figure are reported in Figure 1 – source data 1.

**Figure 4 – figure supplement 3:**
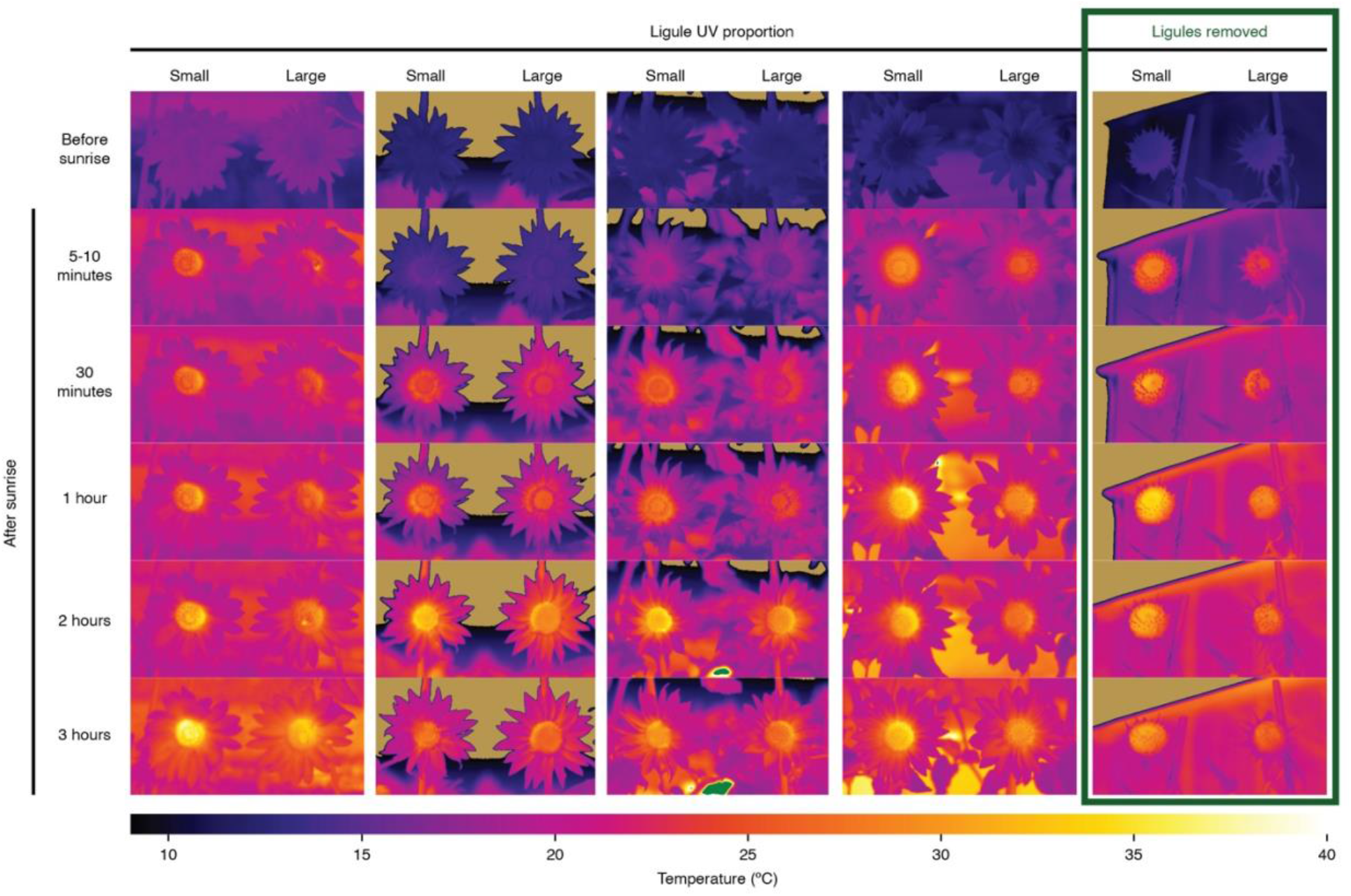
Inflorescence temperature time series. Infrared images of East-facing inflorescences of sunflowers with large (LUVp = 1) or small (LUVp < 0.15) floral UV patterns taken in the summer of 2017. No additional difference was observed in pictures taken more than three hours after sunrise. While no difference in temperature was observed in ligules, the centre (disk) of the inflorescence was consistently warmer in plants with small LUVp. However, this effect is independent of ligule UV patterns, since it persists in inflorescences in which ligules were removed (right-most column). Pollinator visits were severely reduced for inflorescences with ligules removed. Bumblebees can be seen on the disk of inflorescences with large LUVp in the leftmost column of pictures, at 5-10 minutes and 2 hours. Temperatures values outside of the 10-40 °C interval are shown in beige (<10 °C) or green (>40 °C).

**Figure 4 – figure supplement 4:**
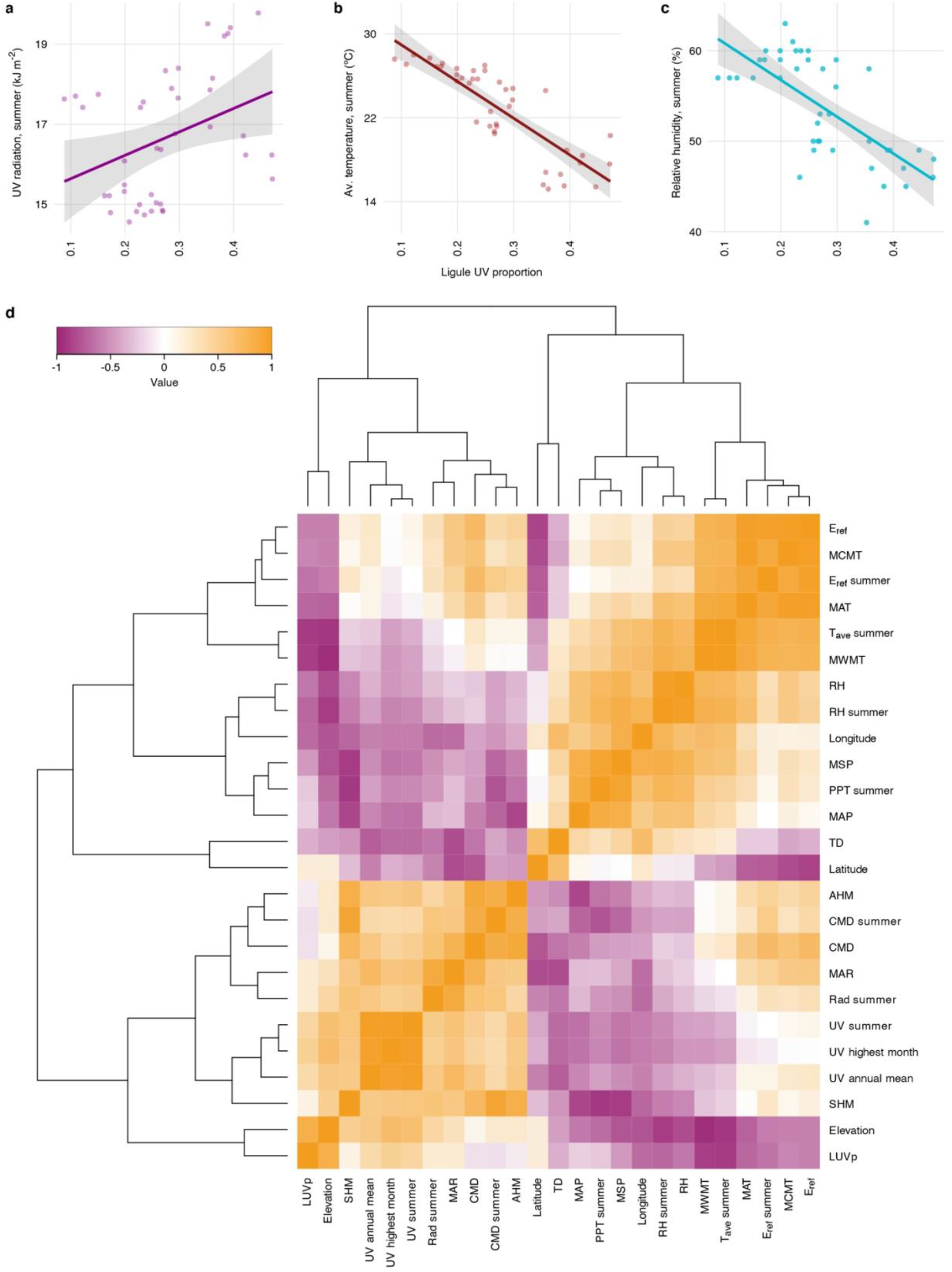
Correlations between LUVp and environmental variables in *H. petiolaris*. **a,** Correlation between average LUVp for different populations of *H. petiolaris* and summer UV radiation (*R^2^* = 0.11, *P* = 0.02), **b,** summer average temperature (*R^2^* = 0.69, *P* = 10^-11^), or **c,** summer relative humidity (*R^2^* = 0.47, *P* =4.4 × 10^-7^). Grey areas represent the 95% confidence interval. **d,** Spearman correlation heatmap for LUVp and environmental variables. Source data for this figure are reported in Figure 1 – source data 1.

## Source data

**Figure 1 - source data 1 (Figure 1 - source data 1.xlsx):** Population used in this study, average LUVp values and environmental variables.

**Figure 1 - source data 2 (Figure 1 - source data 2.xlsx):** Individuals used in this study, LUVp values and Chr15_LUVp SNP genotypes.

**Figure 2 - source data 1 (Figure 2 - source data 1.xlsx):** LUVp values and Chr15_LUVp SNP genotypes for F_2_.

**Figure 3 - source data 1 (Figure 3 - source data 1.xlsx):** Flavonols in methanolic extractions of sunflower ligules and Arabidopsis petals.

**Figure 4 - source data 1 (Figure 4 - source data 1.xlsx):** Pollinator experiment data.

**Figure 4 - source data 2 (Figure 4 - source data 2.xlsx):** Ligules and leaves desiccation experiment data.

**Figure 4 - source data 3 (Figure 4 - source data 3.xlsx):** GEA results for the HaMYB111 region.

## Notes

### Competing Interest Statement

The authors have declared no competing interest.

